# Effect of two activators on the gating of a K_2P_ channel

**DOI:** 10.1101/2024.04.19.590249

**Authors:** Edward Mendez-Otalvaro, Wojciech Kopec, Bert L. de Groot

## Abstract

TREK1, a two-pore-domain (2P) mammalian potassium (K^+^) channel, regulates the resting potential across cell membranes, presenting a promising therapeutic target for neuropathy treatment. The gating of this channel converges in the conformation of the narrowest part of the pore: the selectivity filter (SF). Various hypotheses explain TREK1 gating modulation, including the dynamics of loops connecting the SF with transmembrane helices and the stability of hydrogen bond (HB) networks adjacent to the SF. Recently, two small molecules (Q6F and Q5F) were reported as activators to affect TREK1 by increasing its open probability in single-channel current measurements. Here, using molecular dynamics (MD) simulations, we investigate the effect of these ligands on the previously proposed modulation mechanisms of TREK1 gating compared to the apo channel. Our findings reveal that loop dynamics at the upper region of the SF exhibit only a weak correlation with permeation/ non-permeation events, whereas the HB network behind the SF appears more correlated. These non-permeation events arise from both distinct mechanisms: A C-type inactivation (resulting from dilation at the top of the SF), which has been described previously; and a carbonyl flipping in a SF binding site. We find that, besides the prevention of C-type inactivation in the channel, the ligands increase the probability of permeation by modulating the dynamics of the carbonyl flipping, influenced by a threonine residue at the bottom of the SF. These results offer insights for rational ligand design to optimize the gating modulation of TREK1 and related K^+^ channels.

## Introduction

The two-pore domain (K_2P_) channels are a family of potassium (K^+^) channels found in mammalian cells, primarily responsible for generating leak currents that regulate the negative resting potential across the cell membrane [1, 2, 3, 4]. Modulation of this hyperpolarization is not passive but rather influenced by factors including voltage [5], pH [6], temperature [7, 8], lipids [9, 10, 11], membrane stretch [12, 13, 14], and both endogenous and exogenous ligands [15, 16, 17]. Structurally, K_2P_ channels consist of two subunits, each with four transmembrane helices (TM1 to TM4), two pore domains (P1 and P2), and a selectivity filter (SF) strand in each pore domain (SF1 and SF2). Upon assembly of both subunits, the channel forms a pseudo-tetramer. K_2P_ channels also possess a unique extracellular domain known as CAP, located at the top of the SF [2, 3].

Among the K_2P_ family, the TWIK-related potassium channel 1 (TREK1) is of particular interest as a potential therapeutic target for various disorders including cardiac arrhythmia, stroke, depression, and epilepsy [17, 18, 19]. Modulation of TREK1 currents can influence action potential thresholds, making it significant for developing new anesthetics and analgesics [20, 17]. Efforts to elucidate the TREK1 mechanism include X-ray structures of the truncated channel resolved under different potassium concentrations [21], in the presence of PIP_2_ [22], and co-crystallized with ligands that can lock ion passage [23], enhance the open probability (P_o_), or alter the channel current amplitude [22, 21, 24]. Cryo-EM densities of TREK1 in the presence of anionic and zwitterionic lipids, causing channel opening and blocking respectively, have also been reported [25].

TREK1 exhibits structural asymmetries between each pore domain per subunit (Figure 1 A-F). The P1 domain includes a CAP extracellular domain, and its SF1 strand contains an asparagine (N147) at the top of the filter beyond the S0 binding site. Additionally, the P1 helix contains a phenylalanine (F134) which flanks N147. Conversely, the P2 helix lacks the CAP domain and possesses an aspartate (D256) at the equivalent position of SF1, which is flanked by a tyrosine (Y234) rather than a phenylalanine. Moreover, the loop that connects SF2 with the TM4 helix (denoted SF2-TM4 loop) is longer than its SF1-TM2 counterpart. Notably, TREK1’s selectivity filter primary sequence TIGFG differs from the common TVGYG signature sequence, possessing an isoleucine and phenylalanine instead of valine and tyrosine [26, 16, 21].

**Figure 1.**
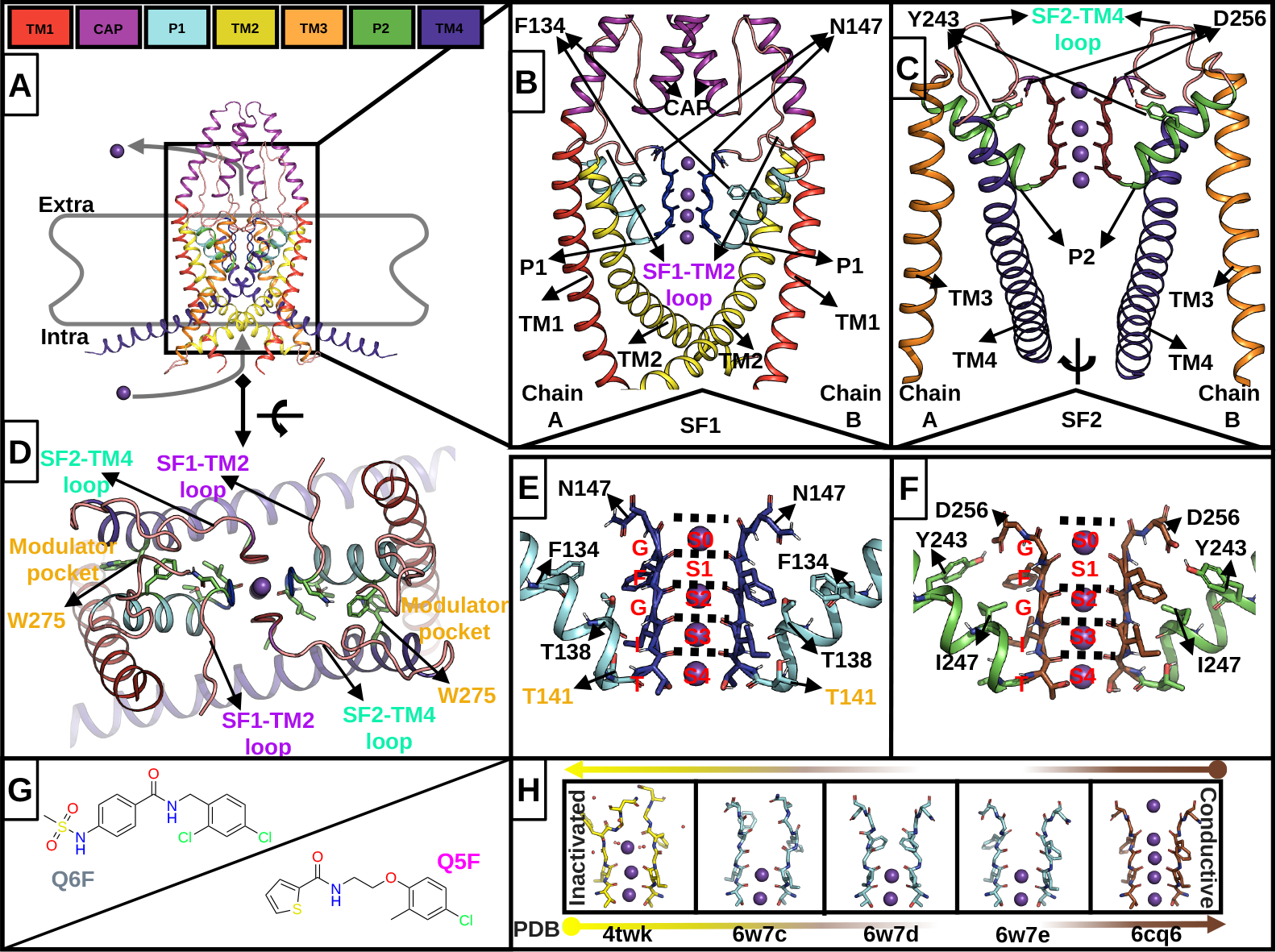
: **A**. TREK1 is a homodimer, and each subunit has two asymmetric pore domains. **B**. Pore domain 1: Notice the CAP domain and the loops that link CAP with the P1 helix, and the SF1 with the TM2 helix. **C**. Pore domain 2: Notice the long loop that links the SF2 with the TM4 helix. **D**. Upper view of TREK1: The CAP and P2 helices were removed for clarity. We highlight the lateral groove formed by the TM4 and P1 helices interface, called the “modulator pocket”. **E**. The SF1 and the P1 helix: Residues of interest are highlighted. **F**. The SF2 and the P2 helix: The equivalent residue position of SF1 is also highlighted here. **G**. Ligands ML335 (Q6F) and ML402 (Q5F) that increase the open probability in TREK1. **H**. SF1 mainchain conformations from representative PDB structures: 4twk represents a putative C-type inactivated SF1 state; notice the co-crystallized water molecules. 6w7c, 6w7d, and 6w7e represent TREK1 SF1 structures at 1, 10, and 30 mM K^+^, respectively, while 6cq6 represents the TREK1 SF1 structure in the apo canonical state.

Similar to other K_2P_ channels, TREK1’s TM4 helix can adopt either an “up” or a “down” state. The “up” state involves TM4 rotating and shifting upward toward TM2, whereas the “down” state positions TM4 nearly 45^*°*^ with respect to the membrane [25, 27]. Recent Cryo-EM data indicates an asymmetry in the “up”/”down” equilibrium between the subunits [25]. Previous evidence suggests that the “up” state is more conductive than the “down” state, with a mechanism that is likely independent of other potential gates [28, 29, 30, 31, 32]. The lack of a classical “helix bundle crossing” or lower gate in TREK1 [33, 34], and the fact that the “up /down” states do not appear to fully impede permeation [29, 35, 36], suggests a gating mechanism that is primarily driven at the SF level.

One proposed mechanism of gating in TREK1 involves asymmetric order-disorder transitions in the SF structure: the absence of ions in S1 and S2 triggers pinching in SF1 and a dilation in SF2, that disrupts the canonical conductive geometry and impedes channel permeation, resembling a C-type inactivation [21] (Figure 1 H). This mechanism correlates with the pharmacological activity of two molecules, ML335 (Q6F) and ML402 (Q5F) (Figure 1 G), which enhance the open probability of TREK1 [22], by modulating a specific gate affecting the inactivation process. The transduction of external cues into the gating mechanism of TREK1 remains elusive, and some possible three mechanisms are summarized in Table 1 (Also Figure 1, B-E, and Figure S7 A-C).

**Table 1:** Possible gating mechanisms converging in the SF.

| Channel region | Hypothesis | Details |
| --- | --- | --- |
| **I) SF2-TM4 loop** | Loop stiffness | Mutagenesis studies indicate that cues such as temperature and pressure converge in this region [8]. The interaction between G260 (SF2-TM4 loop), Y270 (TM4), and E234 (TM3) transduces these signals into the channel [21]. |
| **II) SF1-TM2 loop** | Loop stiffness | The I148T mutation increases the I and $P_o$ of the channel [8]. Mutations in charged residues (H, E, R) at the end of this loop modify the channel's activity [37, 38]. |
| **III) P1 & TM4 helix** | HB network changes behind SF1 | Simulations suggest that a HB between the $N_\epsilon$ of W275 (TM4) and OH of T141 (P1) destabilizes the backbone of residue I143, which forms the S3 in the SF1 [39]. Conversely, experiments indicate that W275 can open TREK1 when pushed inward the SF by lipids [25]. |

We hypothesize that insights into channel gating could be resolved by investigating the effects of Q6F and Q5F ligands on previously proposed gating mechanisms (Table 1) for TREK1 behavior and to this end, we performed extensive MD simulations on both the apo and ligand-bound channels. Importantly, we find that the stiffness of the SF1-TM2 and SF2-TM4 loops (Hypothesis I and II) are unlikely to modulate the gating in TREK1, as their dynamics do not correlate with channel permeation /non-permeation events. On the other hand, we find that the hypothesis III involving the residue T141 is more likely, as its side-chain (SD) dynamics directly relate to non-permeation events in the apo channel, suppressed in the ligand-bound state.

Interestingly, we observe that the non-permeating regimes arise from two distinct mechanisms: 1.) A C-type-like inactivation, which involves disorder transitions at the S0 and S1 binding sites, and has been previously described in TREK1 by Lolicato *et al*. [21] 2.) The flipping of a carbonyl at the S3 site in the SF, which is prevented by ligands to a greater extent, and is directly related to the T141 side-chain dynamics.

## Materials and Methods

### Computational models

Initialize structures of the TREK1 channel in its apo state (PDB: 6cq6) and bound to Q6F (PDB: 6cq8) and Q5F (PDB: 6cq9) [22] were taken from the Protein Data Bank (PDB). These structures miss a loop in the subunit A (residue 113 to 124). The loop was manually modeled by taking the same loop coordinates that are resolved in the subunit B. The same strategy was used for the terminal end in the subunit B (residue 317 to 321), which is fully resolved in the subunit A. The structures were protonated according to their standard protonation state at pH 7 and were inserted into a POPC membrane (1-palmitoyl-2-oleoyl-phosphatidylcholine) that was surrounded by water molecules and K^+^ and Cl^-^ ions, resulting in a hexagonal box with a K^+^ concentration of *≈* 500 mM, using the CHARMM-GUI web server [40, 41, 42, 43]. The protein termini were capped with ACE and NME.

### Simulation setup

All simulations were performed with the MD software GROMACS 2021.7 [44, 45, 46, 47, 48, 49]. We used two different force-fields: CHARMM36m [50] (with CHARMM TIP3P water [51], CHARMM36 lipids [52, 53], CgenFF ligands [54, 55, 56, 57], and the Beglov & Roux ions [58]) and Amber14SB [59] (with TIP3P water [60], Slipids [61, 62], GAFF2 ligands [63, 64, 65], and the Joung & Cheatham [66] ions). Systems were equilibrated in six steps using the default scripts provided by CHARMM-GUI with GROMACS 2021.7, with a final equilibration step without restrains of 50 ns. For the CHARMM systems, ligand parameters were obtained from the CgenFF server (https://cgenff.silcsbio.com/) and were converted to a GROMACS-compatible format using the cgenff_charmm2gmx.py script (http://mackerell.umaryland.edu/charmm_ff.shtml#gromacs); for the Amber systems, GAFF2 parameters were generated using the Antechamber Python parser interface (ACPYPE) [67].

Production simulations were performed with the leapfrog integrator and a 2 fs time step. Temperature was maintained at 323.15 K using a Nosé-Hoover [68] thermostat and pressure was maintained at 1 bar using a semi-isotropic Parinello-Rahman [69] barostat. All hydrogen bonds were constrained using the LINCS [70] algorithm. For CHARMM36m, Van der Waals interactions were force-switched off between 1.0 and 1.2 nm and Coulomb interactions were computed using the PME method [71] with a 1.2 nm real-space cutoff. For Amber14SB the interaction cutoff was 1.0 nm, a dispersion correction for energy and pressure was applied and the PME cutoff was 1.0 nm. Each system was simulated with an external electric field along the Z-axis, generating a membrane voltage of *≈* 300 mV. The voltage was calculated as:

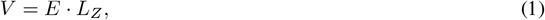

where *E* is the applied field and *L*_*Z*_ is the length of the simulation box along the Z-axis [72, 73]. Frames were saved every 40 ps and 10 replicates of 1 *µ*s each were carried out for each system (apo, Q5F, and Q6F) and force field yielding a total simulation time of 60 *µ*s.

### Analysis

For all analyses of distances, atom positions, angles and mean square displacements, we used Python 3 [74] scripts and the Numpy [75] and MDAnalysis [76, 77] libraries. The results were plotted using the Matplotlib [78] library in Python Jupyter Notebooks.

Permeation events were identified using a custom script (available at https://github.com/huichenggong/Sfilter_Cylinder) and the the conductive current was calculated by fitting *f* (*x*) = *Cx* + *b* to the cumulative permeation traces of the conductive channel (i.e., when the channel is actively permeating ions). In this work, we define a channel to be non-permeating if 30 ns have elapsed since a permeation event has occurred. This time was chosen based on the distribution of all permeation times computed from the simulations (Figure S7 F, E). We also compute the average current (i.e., total permeation events divided by simulation length) that includes both conductive and non-conductive states and is lower than the conductive current. The permeation probability was calculated as the fraction of time in which the filter does not conduct ions with respect to the total simulation time.

Results showed are averaged over the 10 independent replicates and error bars represent the standard error of the mean. For the PCA (principal component analysis) of the solved structures, we took the TREK1 coordinates reported by Lolicato *et al*. [22, 21], Pope *et al*. [23], Schmidpeter *et al*. [25], and Pike *et al*. [79]. We retrieved the structures of the PDB database and extracted the SF1 coordinates, then we concatenated them in an ensemble. We calculated the eigenvalues and eigenvectors of the covariance matrix of this ensemble of experimental structures after rotational and translational fitting to the SF1 of the apo conductive structure (6cq6). Finally, we projected our whole sampling into the first two eigenvectors (these eigenvectors describe *≈* 77% of the variance). As an order parameter for the SF1 state, we used the weighted average of these first two eigenvectors 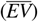:

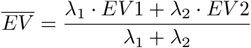

where *EV1* and *EV2* represent the projection of the simulation onto their respective eigenvectors, and *λ*_1_ and *λ*_2_ denote the variances explained by each of the eigenvectors.

The distribution of water molecules along the Z axis was calculated after dumping the water positions using a custom Fortran script (available in previous work of Kopec *et al*. [80] https://www.nature.com/articles/ s41467-019-13227-w#Sec13). The probability density function (PDF) plots were calculated after properly fitting our channel simulations to the SF of the apo conductive structure (6cq6), and dumping the K^+^ positions along the Z-axis. Then, we converted such positions into densities through a Gaussian kernel estimator using the SciPy [81] library. In this case, the thick line represents the average density, and the shaded region is the confidence interval of the mean.

The interaction fingerprints of the ligands with TREK1 were calculated using ProLIF [82]. All residues in contact with the ligands within a radius of 2 nm were included. The similarity matrix between these fingerprints was computed using the Tanimoto index. In order to identify common interaction modes across the entire sampling time, the similarity matrix was subject to spectral clustering in scikit-learn [83]. All the representative snapshots of the MD simulations were taken using PyMOL [84] and the visualization of the trajectories was done using VMD [85].

## Results

Through simulations employing the Amber force field, we observed minor distortions in the filter and stabilization of residue side-chain dynamics. These findings align with previous research on the MthK K^+^ channel [86] and the prokaryotic KcsA K^+^ channel [87]. As there was no significant difference in the permeation probability of TREK1 with or without activators under this force field (see Figures S12 and S13), our primary focus now turns to the results obtained using the CHARMM force field.

### Ligands increase the permeation probability in TREK1 and decrease the probability of the S3 carbonyl flip

To assess the impact of the ligands on the filter geometry, we projected the simulated trajectories onto the first two eigenvectors resolved from a principal component analysis (PCA) of the SF1 mainchain atoms taken from multiple experimentally resolved TREK1 structures. The first two components were combined into a single parameter 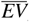 (see methods) that primarily describes distortions at the top of the filter.

We observed a clear separation between non-canonical conformations and those corresponding to the channel in its conductive canonical conformation (Figure 2 B). In the presence of Q6F, the channel is more likely to sample SF states near the conductive conformation, whereas the apo channel and the Q5F-bound channel sample a larger conformational space. We identified one minimum sampled predominantly by the apo channel (*EV1≈* -1.4, *EV2≈* 0.2), that is less frequently sampled by the ligand-bound channel, indicating that both molecules have an effect on filter geometry. Further characterizing this minimum apo-sampled state revealed it to correspond to an SF1 configuration with water bound at S0 and S1, flipped carbonyls at S0 and S1, and a slight distortion in the side chain of F145 (S1) and the carbonyl of S3.

**Figure 2.**
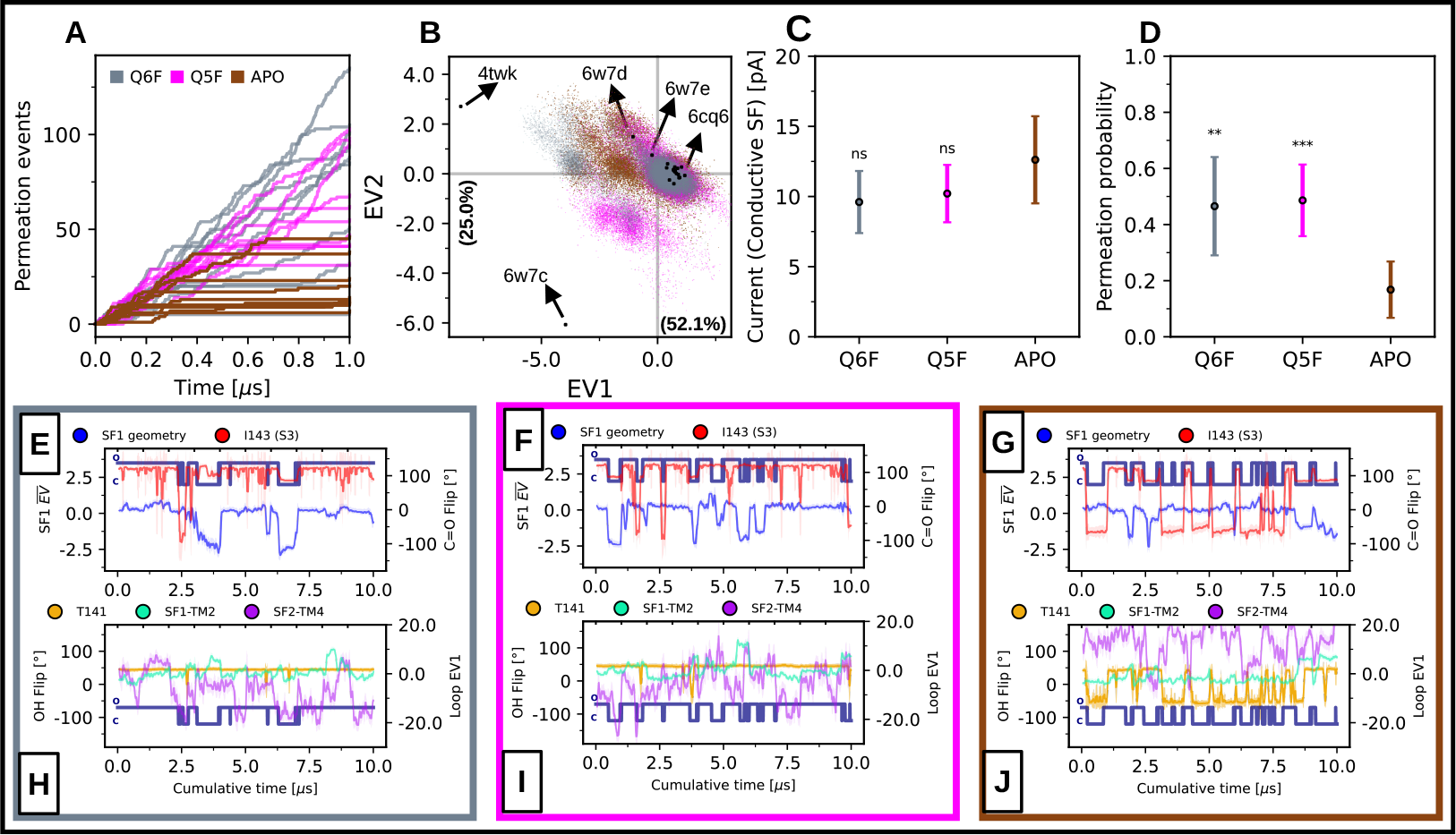
: **A**. Cumulative permeation events over 1 µs (N=10 for each condition). **B**. Total sampling of the holo and apo simulations projected on eigenvectors 1 and 2 of the SF1 main-chain ensemble of experimental structures (30 µs of sampling are projected. EV means eigenvector). **D, C**. Permeation probability and channel current when the SF is conductive (See Methods). Both agree qualitatively with that observed in single trace currents experiments (two-sided T-test for the means. ns: P *>* 0.05. *: P *≤* 0.05. **: P *≤* 0.01. ***: P *≤* 0.001. Error bars correspond to confidence intervals of the mean). **E, F, G**. Two gating mechanisms as a function of cumulative time (N=10): C-type inactivation distortions (given by the weighted average 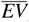); and the carbonyl flip in S3 (given by the N-Cα-C-O dihedral angle in residue I143). The blue-navy traces indicate when the channel is permeating (O) or not permeating (C) ions, respectively. Each panel corresponds to the two holo cases and the apo case, respectively. **H, I, J)** Three hypotheses about the modulation of TREK1 gating by ligands: Dynamics of the SF1-TM2 and SF2-TM4 loops given by the EV1 of their main-chain; and the stability of the HB network behind SF1 given by the χ1 angle of the T141 residue.

With respect to ion permeation, we observed a higher average current in the presence of ligands (Figure S7 A) as well as an increased probability of permeation: the apo channel exhibits prolonged periods of no permeation through the filter (Figure 2 A, D) compared with the ligand-bound case. When calculating the conductive current, we observed no significant difference for the apo or the ligand-bound states (Figure 2 C). This can be attributed to prolonged stretches of low conductance in the apo channel, leading to an overall decreased probability of permeation (Figure 2 A). For conductive stretches, we see no significant difference in current between apo and ligand-bound cases (Figure 2 C). These results are in agreement with previous TREK1 single-channel current measurements [21].

Given the observed slight distortion in the carbonyl of S3 (also observed previously by Lolicato *et al*. [21]), and the fact that previous work suggests that flipping of this group can regulate permeation in TREK2 [28], we analyzed the relationship between permeation/non-permeation events, the S3 backbone dihedral angle (i.e., N-C*α*-C-O), and 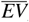 (which primarily describes large distortions at the top of the filter). Importantly, the carbonyl flip in S3 is predominantly observed in the apo channel; whereas in the ligand-bound channel, non-permeation occurs due to distortions in the filter as described by 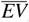 (Figure 2 E, F, G). This change in the S3 dihedral angle suggests that, besides the previously proposed prevention of C-type distortions by the ligands [21], the S3 flipping mechanism is also regulated to a greater extent.

Thus in addition to the prevention of C-type inactivation, ligands largely modulate carbonyl flipping at S3, but it is worthwhile to analyze the mechanism by which C-type inactivation is prevented. Projections along *EV1* revealed that the Q6F-bound channel more frequently samples configurations in the low distortion regime between -0.24 *< EV1 <* 1.17 when compared to apo channel (i.e., *≈* 10% more), consistent with the increased average conductance (Figure S11). Further analysis of the Q6F-bound channel revealed this ligand to hinder the sampling of non-canonical states in the side chains of residues N147 (located at the top of the filter) and F134 (helix P1) (Figure S8). The side chain of residue N147 points canonically outward toward residue F134, which is similar to the D80-W67 interaction pair in the prokaryotic KcsA channel, and the D447-F434 in the *Shaker* C-type inactivated W434F mutant channel. Notably, these channels both display a similar geometry to the distorted SF1 states (Figure S6) observed here. The unique geometry of Q6F (e.g, sulfonamide moiety and longer length) allow for more favorable residue interactions as compared with Q5F (Figure S4, S5 and S10), which in turn promotes the conservation of the canonical geometry at the top of the SF.

### Little evidence to support hypothesis I and II: Ligands reduce loop fluctuations, but with poor correlation to permeation probability

After assessing the effect of the molecules on both the channel filter geometry and the transport function, we evaluated hypotheses I and II regarding the stiffening of the SF1-TM2 and SF2-TM4 loops. We calculated the mean fluctuations of Cα in both loops for the apo and the ligand-bound channels and found no significant difference in the stiffness of the SF1-TM2 loop. However, the SF2-TM4 loop exhibits reduced fluctuations in the middle portion (residues V258-G261) in the presence of Q6F (Figure S1 A, B). To test whether this effect relates to gate modulation, we monitored loop dynamics using the first eigenvector of the main chain for each loop as an order parameter. We observed no change in the SF1-TM2 loop dynamics between conditions, nor any correlation with non-permeation events. Similarly, for the SF2-TM4 loop, although both ligands induce an average shift in the loop’s sampled space, their dynamics are not related to non-permeation events (Figure 2 H, I, J). Hence, we conclude that these two hypotheses regarding the modulation of the channel gating by the ligands are not supported by the data.

### Hypothesis III is most related to the SF non-conducting events: Ligands reduce the flipping of carbonyl S3 by stabilizing the T141 side chain

To test hypothesis III, we measured the χ1 angle of the T141 residue at the bottom of SF1 (end of P1 helix) and the angle formed between: the center of mass of the W275 side chain (located in TM4 helix), its C*γ* atom, and the center of mass of the S4 binding site. Although both ligands induced a shift in the W275 side chain towards more acute angles, the correlation between the W275 dynamics and channel permeation state (Figure S1 E, F, G) is weak. Importantly, T141 dynamics is highly related to non-permeation events in the apo channel, fluctuating between 50^*°*^ and *−*50^*°*^ depending on the dynamics driven by the flip mechanism in S3. In the ligand bound channel, T141 is restricted to an “up” state, which directs the hydroxyl towards the SF1 backbone, creating an anchor for hydrogen bond formation with the backbone amide of I143, which in turns hinders the S3 flipping process (100^*°*^ *→ −*50^*°*^) and prevents non-permeation regimes (Figure 2 H, I, J).

## Discussion

TREK1 is modulated by various external cues such as membrane stretch, temperature, intra and extracellular pH, as well as exogenous and endogenous ligands. Understanding how these signals are transduced into K^+^ permeation, and ultimately, the resting potential, holds significant pharmaceutical importance.

Previous studies suggest that TREK1 gating primarily converges at the SF, where pinching and dilation of the S2 & S1 binding sites regulate the gate [21]. Additionally, we find that, besides this C-type inactivation, TREK1 gating involves carbonyl flipping at S3, as previously proposed for TREK2 [28], TWIK-1 [88] and hERG [89] K^+^ channels. This flipping can occur spontaneously when S3 is empty, but it primarily occurs in the presence of a water molecule in S3, increased hydration levels in the modulator pocket, and water molecules behind the SF1, which collectively result in a sustained flip (Figure S3). However, complete stabilization of the gating mechanism only occurs when the T141 side-chain transitions to its horizontal “down” state. A valid question is whether there is a coupling between the two gates (C-type inactivation and the carbonyl flip in S3). Our evidence indicates that such coupling is weak. Nevertheless, in the case of the ligand-bound channel, the population of states sampled from both the C-type-inactivated channel and the channel with the flipped carbonyl is much smaller than in the apo case (Figure S11).

We find the ligands to exert a variety of effects on the channel, that are linked to the flipping mechanism previously described: 1.) They significantly reduce the probability of a water molecule being present in S3 (Figure S2 A, B, C), 2.) They reduce the energy barrier that the ions have to cross in S3 (Figure S2 A, B, C), 3.) They alter the hydration levels behind the SF1 and around T141 (Figure S9), and 4.) They stabilize T141 in the “up” state, preventing complete permeation blockage (Figure 2).

Regarding the C-type inactivation in the ligand-bound channel, this occurs when the S0 and S1 sites are empty and filled with water molecules. In such a state, ligands adopt a conformation that reduces interactions with aromatic side chains of F145 (S1) and F134 (P1 helix), as well as the N147 side-chain at the top of the SF1 (Figure S3). Here we observe a stronger effect from Q6F that has a sulfonamide group, allowing it to interact more strongly with N147 as compared to the thiophene ring of Q5F (Figures S4, S5). The result is that the Q5F-bound channel more frequently samples C-type inactivated regions compared to Q6F. Taken together, our results suggest that both molecules modulate the carbonyl flip mechanism at S3; however, due to its chemical structure, Q5F exerts a smaller effect on the C-type inactivation mechanism than Q6F (Figures S4, S5, S10).

In both the C-type inactivation mechanism and the carbonyl flip in S3, we find that water plays an important role: its presence in S0 and S1 promotes distortions at the top of the filter and in the phenyl sidechains of F134 (P1) and F145 (S1) (Figure 3 D, Figure S3, Figure S8 A, B, C), altering their hydrophobic packing and inciting the C-type inactivation. Meanwhile, the presence of water in S3 and around residue T141 enhances the carbonyl flip and the “down” state of the hydroxyl in T141 (Figure 3 D, Figure S2, Figure S3, Figure S9). Therefore, we hypothesized that the degree of hydration surrounding the filter is a key element for modulation in TREK1 and that the ligands are modifying it directly.

**Figure 3.**
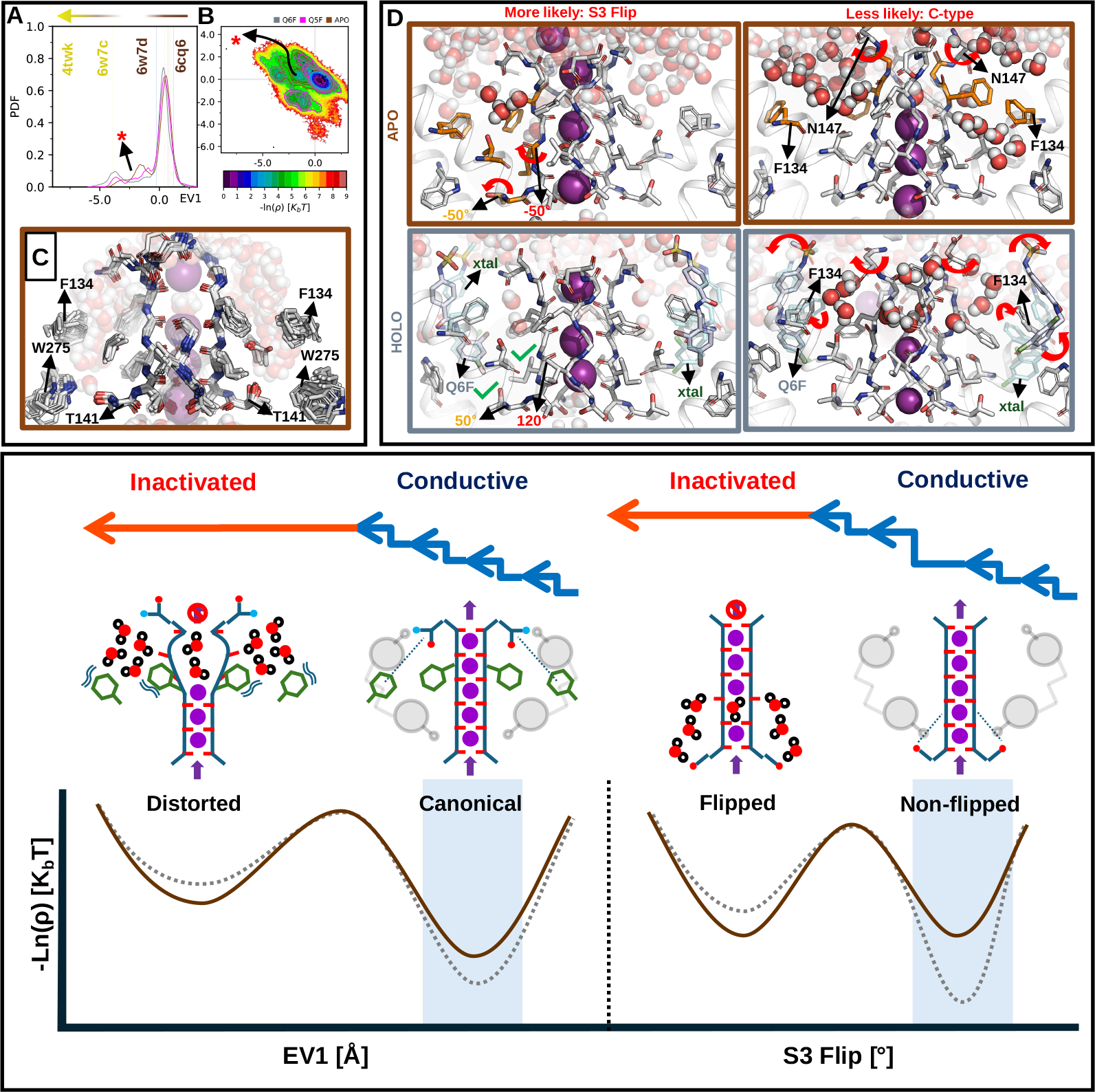
: **A**. Probability density function (PDF) of the first eigenvector (EV1) describing the state of SF1. **B**. Negative logarithm of the PDF (“Free Energy”) of EV1 & EV2 describing the state of SF1. The red asterisk indicates the one minimum mainly sampled by the apo-channel. The isocontours indicate which regions are sampled by the channel depending on whether it is bound to ligands (Q6F, Q5F) or not (apo). **C**. Representative snapshots of the conformations sampled at the minimum indicated by the red asterisk in A and B. Note the presence of water molecules (transparent space filling model) in S1 and S0, and the C-type-like distortions. **D**. Two mechanisms of TREK1 gating. The ligands modulate mainly the S3 carbonyl flip. The lower panel summarizes the two mechanisms presented above; the dashed case represents presence of the ligand.

The level of hydration around the SF might be related to the difference in the filter primary sequence (TIGFG instead of the common TVGYG sequence). The replacement of a tyrosine by a phenylalanine eliminates a potential hydrogen bond that would stabilize the side chain, replacing it with a hydrophobic interaction with F134 (P1). This hydrophobic interaction has also been proposed as key in the inactivation of the hERG channel [89]. The same might occur with the substitution of valine by isoleucine, making the side chain slightly longer and more hydrophobic. This reasoning has previously been proposed to explain the relationship between filter sequence differences and conductive states in other potassium channels, in particular TWIK-1 [88].

The correlation between the change in hydration levels around the modulator pocket and permeation events is supported by previous structural, functional, and computational studies. For instance: The inactivated structure of TREK1 (PDB: 4twk, Figure S6) exhibits co-crystallized water molecules behind SF1 and inside S0 and S1. The presence of hydrophobic moieties in the modulator pocket (i.e., lipids [90, 25], hydrophobic mutations (e.g., G137I mutation [91]), and aromatic residue packing in the pocket [92]) can activate the channel, likely by decreasing hydration levels in the region, similar to those observed in our simulations (Figure S1 D. S9). Previous simulations by Lolicato *et al*. [21] have also identified changes in water density around SF1 during filter distortion, similar to observations in simulations of rapid inactivation in hERG [89] and MthK channels [86].

With respect to ligand design, we hypothesize that a molecule with a charged or more polar upper end that coordinates with the N147 residue located at the top of the SF1 might be even more efficient for prevention of a C-type inactivation. Additionally, the introduction of a double bond in the linker between the upper and lower ends of the ligand might help to stabilize the sulfonamide in a configuration conducive to N147 interactions and to reduce the overall conformational entropic penalty, both potentially improving binding affinity. Further, increasing the ligand length and hydrophobicity at the lower end with an aromatic double ring could enhance the ligand effect by wedging TM4 “up” and preventing fluctuations. Such a modification would also help to preserve the dehydrated state behind the SF by packing more closely against the side chains of T141 and T142 (that form S4), and the hydrophobic patch residues F170 to A175 (in TM2) and V274 to A283 (in TM4). This region is analogous to the region identified in TRAAK by Kopec *et al*., where coupling between the activation gate and the selectivity filter gate was mediated by hydrophobic contacts [80]. It also corresponds to the region of increased hydration observed in the apo channel (Figure S9) and the region with significant ligand interactions (Figure S10). Some of these modifications have already been investigated and a ligand with both a negatively charged upper end and a bulkier lower end [93] binds to the same modulator pocket and enhances channel conductance.

## Conclusion

In this study, we have addressed the effect of two activators on TREK1 gating, focusing particularly on three possible hypotheses. The stiffness of the loops at the top of SF1 (Hypothesis I) and SF2 (Hypothesis II) weakly correlates with non-permeation events overall. Hypothesis III appears more likely because the side-chain dynamics of residue T141 (involved in this hypothesis) are highly related to permeation /non-permeation events.

Furthermore, the non-conductive state of the channel arises from two mechanisms: 1) C-type inactivation at the top of the filter, involving the state of the N147-F134 interacting pair. 2) Carbonyl flipping in the SF1, highly modulated by ligands and directly linked to the dynamics of the side chain of residue T141, as well as the hydration levels behind SF1 and within the modulator pocket.

Therefore, we contribute to understanding the gating mechanism of TREK1 by studying the effects of two activators in detail. Our findings may help to design new and more effective molecules that modulate TREK1 and similar K^+^ channels at the level of their gating mechanisms.

## Author Contributions

E.M.O., W.K. and B.L.G. designed the research. E.M.O. carried out all simulations, and analyzed the data. E.M.O., W.K. and B.L.G. wrote the article.

## Acknowledgments

Edward Mendez-Otalvaro thanks Chenggong Hui, Carter J. Wilson, and Andrei Mironenko for providing some of the scripts to analyze the simulations. Also system administrators Ansgar Esztermann & Martin Fechner for their help in technical support; and all members of the group Computational Biomolecular Dynamics. Acknowledgments to Petra Kellers and Carter J. Wilson for kindly proofreading the manuscript. This research was funded by Leibniz Forschungsinstitut für Molekulare Pharmakologie im Forschungsverbund Berlin e.V. (FMP K305/2020) [Leibniz SAW-2018-FMP-a-P5label (PSBICH6019)].

## Supplementary Material

**Figure S1:**
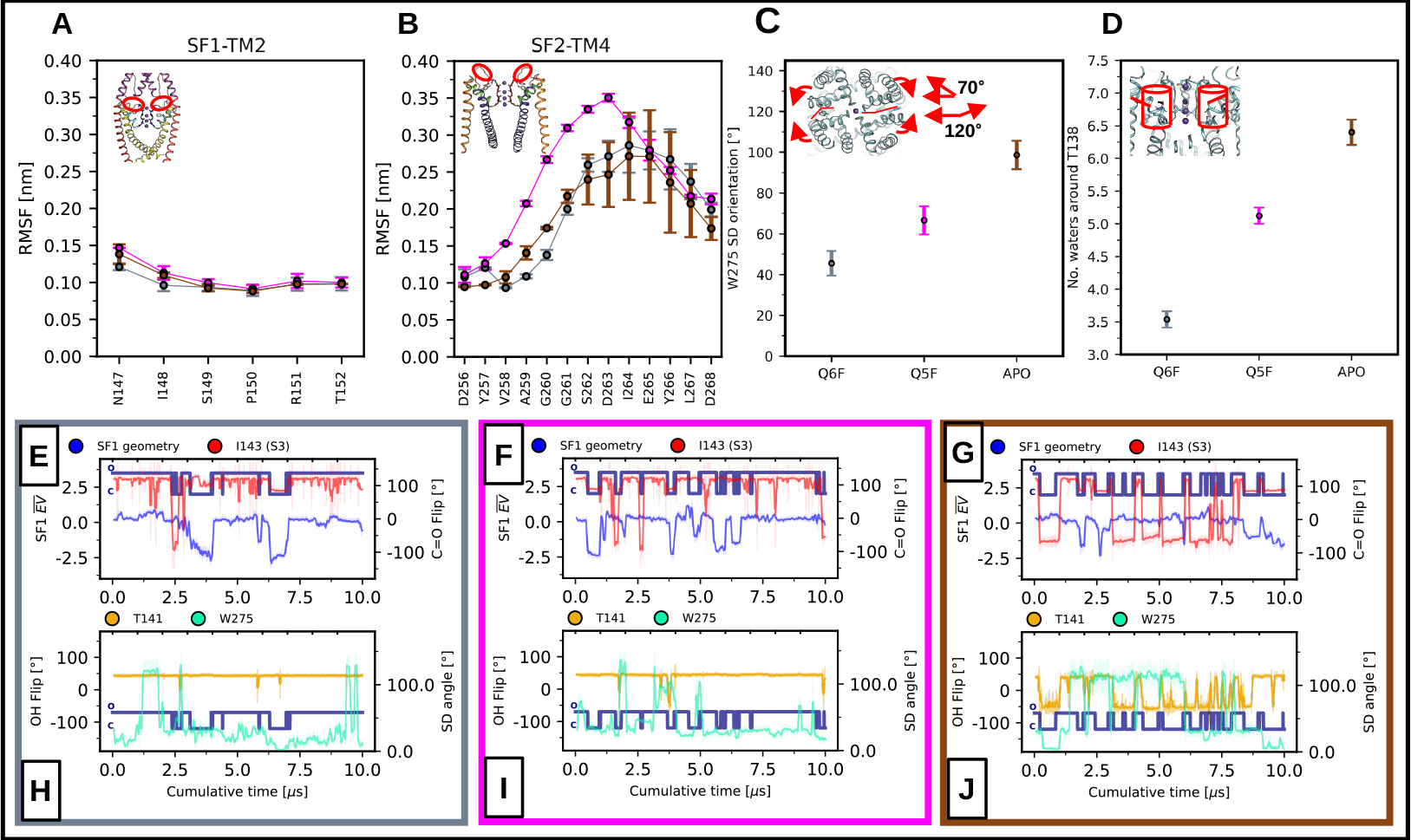
**A, B**. RMSF of Cα of both SF1-TM2 and SF2-TM4 loops. **C**. W275 side-chain orientation with respect to the S4 of the filter. **D**. Number of water molecules around 0.5 nm of T138, which is located behind SF1. **E, F, G**. Two gating mechanisms as a function of cumulative time (N=10): C-type inactivation distortions (given by the weighted average 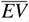); and the carbonyl flip in S3 (given by the N-Cα-C-O dihedral angle in residue I143). The blue-navy traces indicate when the channel is permeating (O) or not permeating (C) ions, respectively. Each panel corresponds to the two holo cases and the apo case, respectively. **H, I, J)** Two order parameters about the modulation of TREK1 gating by ligands: W275 side-chain orientation with respect to the S4 of the filter; and the stability of the HB network behind SF1 given by the χ1 angle of the T141 residue.

**Figure S2:**
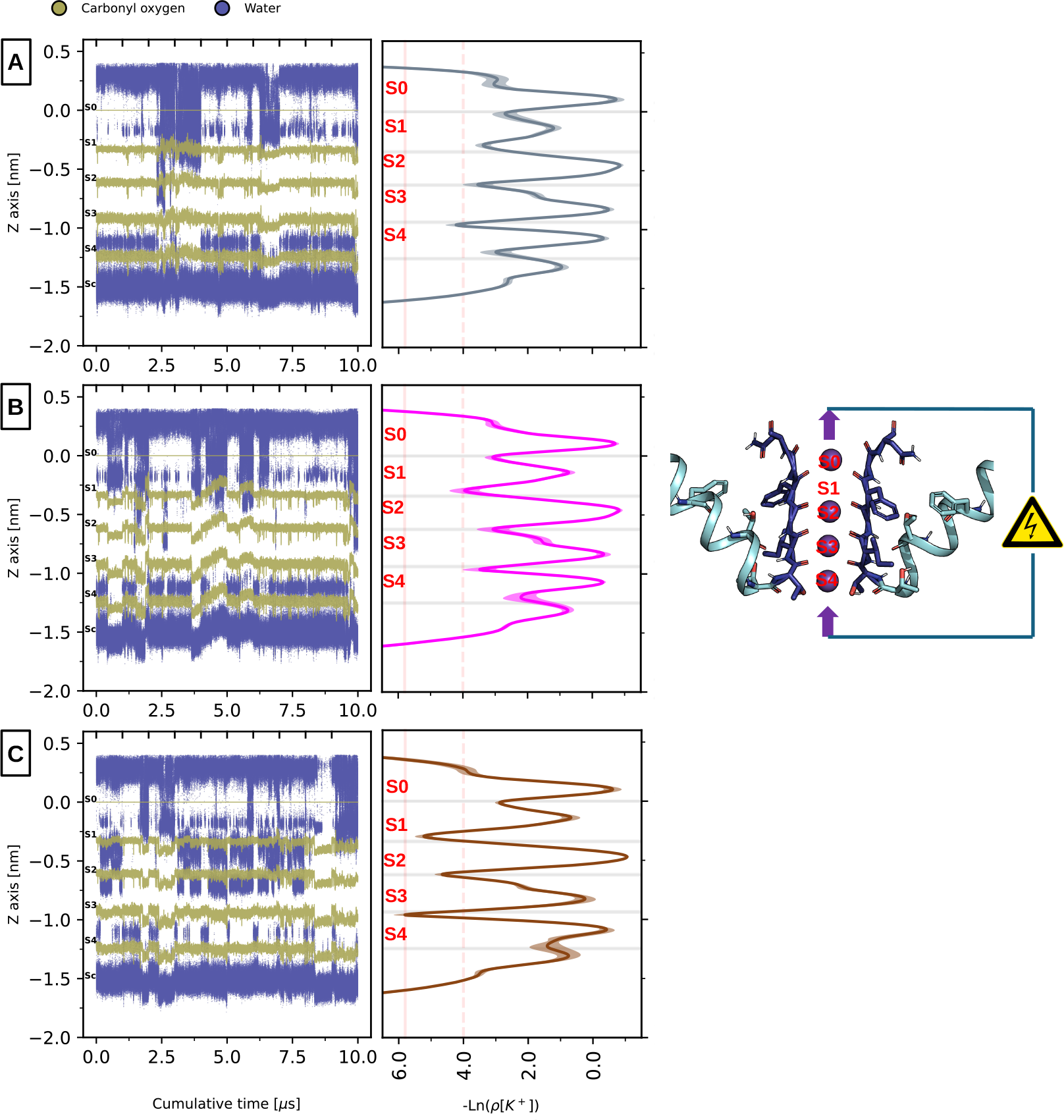
Water positions along the Z-axis of the channel’s filter. The side panels correspond to the negative logarithmic density of K^+^ positions along the Z-axis for Q6F **A**., Q5F **B**., and apo **C**.

**Figure S3:**
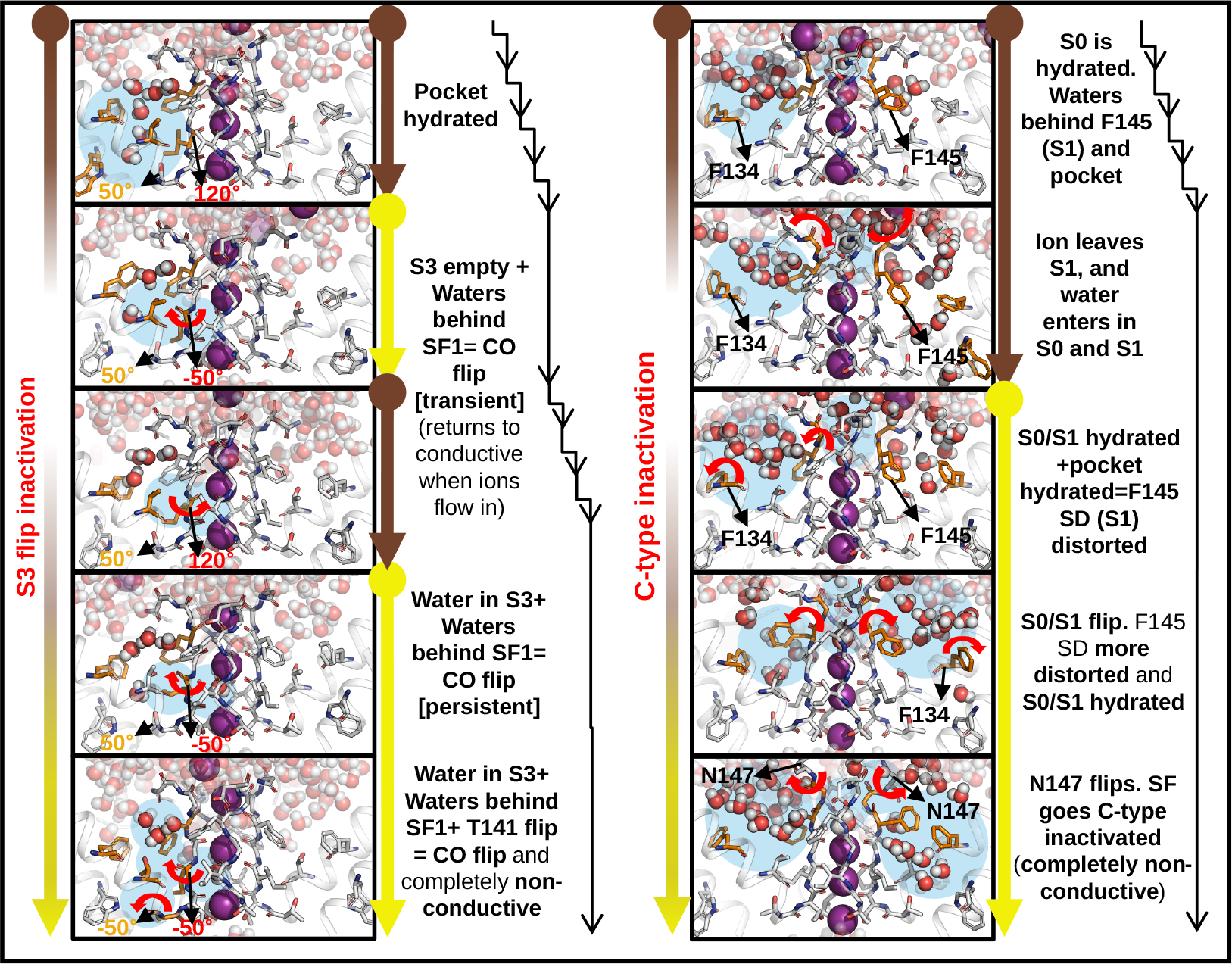
Molecular details of the two mechanisms of gating in TREK1: Carbonyl flipping at S3 and C-type inactivation.

**Figure S4:**
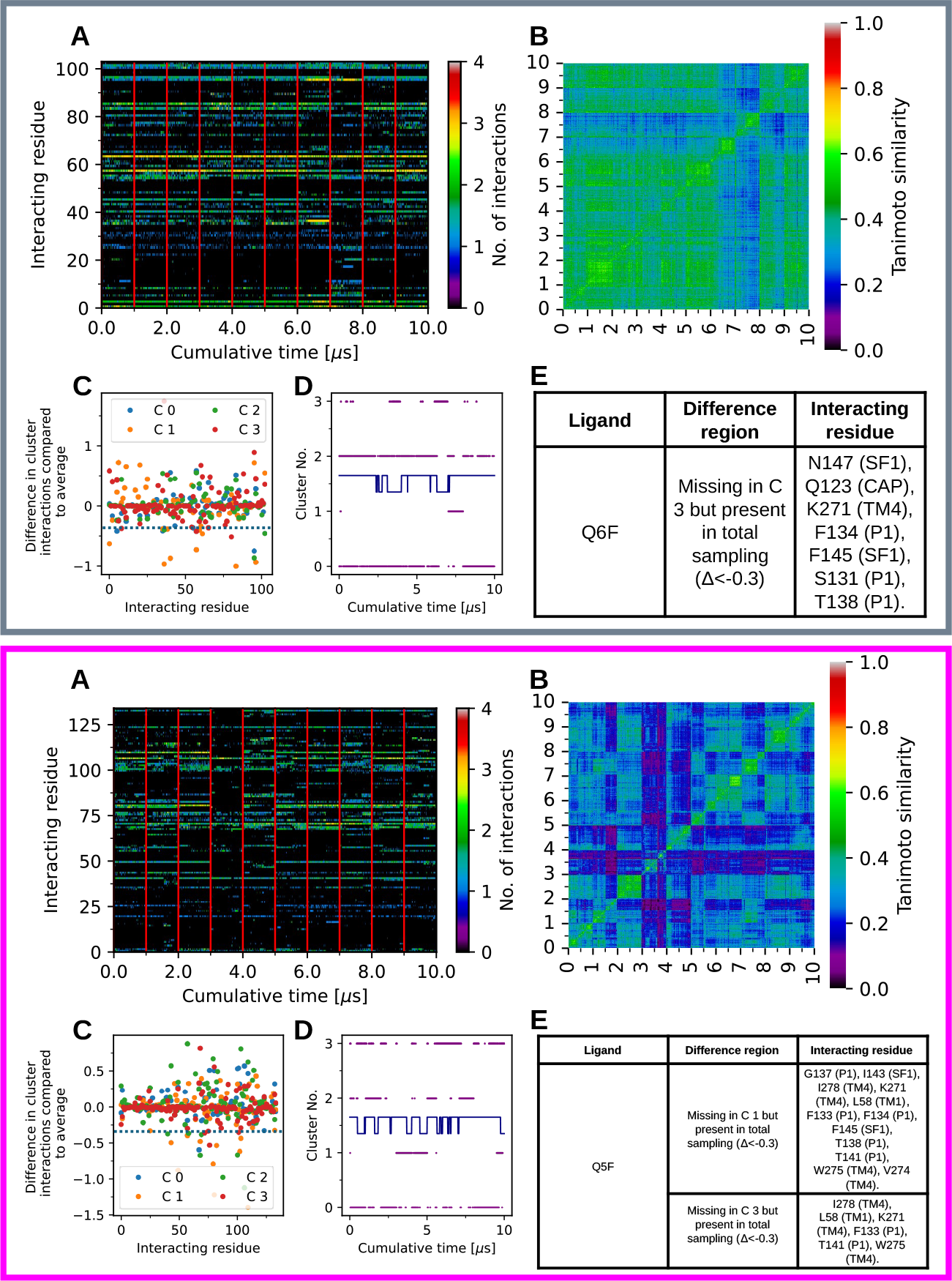
Binding modes for the Q6F **Top panel** and Q5F **Bottom panel** activators. **A**. Interaction profile of the molecule with the protein over cumulative time. **B**. Similarity matrix of this interaction profile. **Cluster analysis of the similarity matrix: C**. shows the difference between the average number of interactions of each cluster with respect to the total interaction profile. **D**. shows each of the clusters as a function of time, with the blue-navy trace indicating whether the channel is permeating or not permeating ions.. **E**. Table with the interactions present in the total interaction profile but absent in the indicated cluster; the limit for evaluating these differences is given in parentheses.

**Figure S5:**
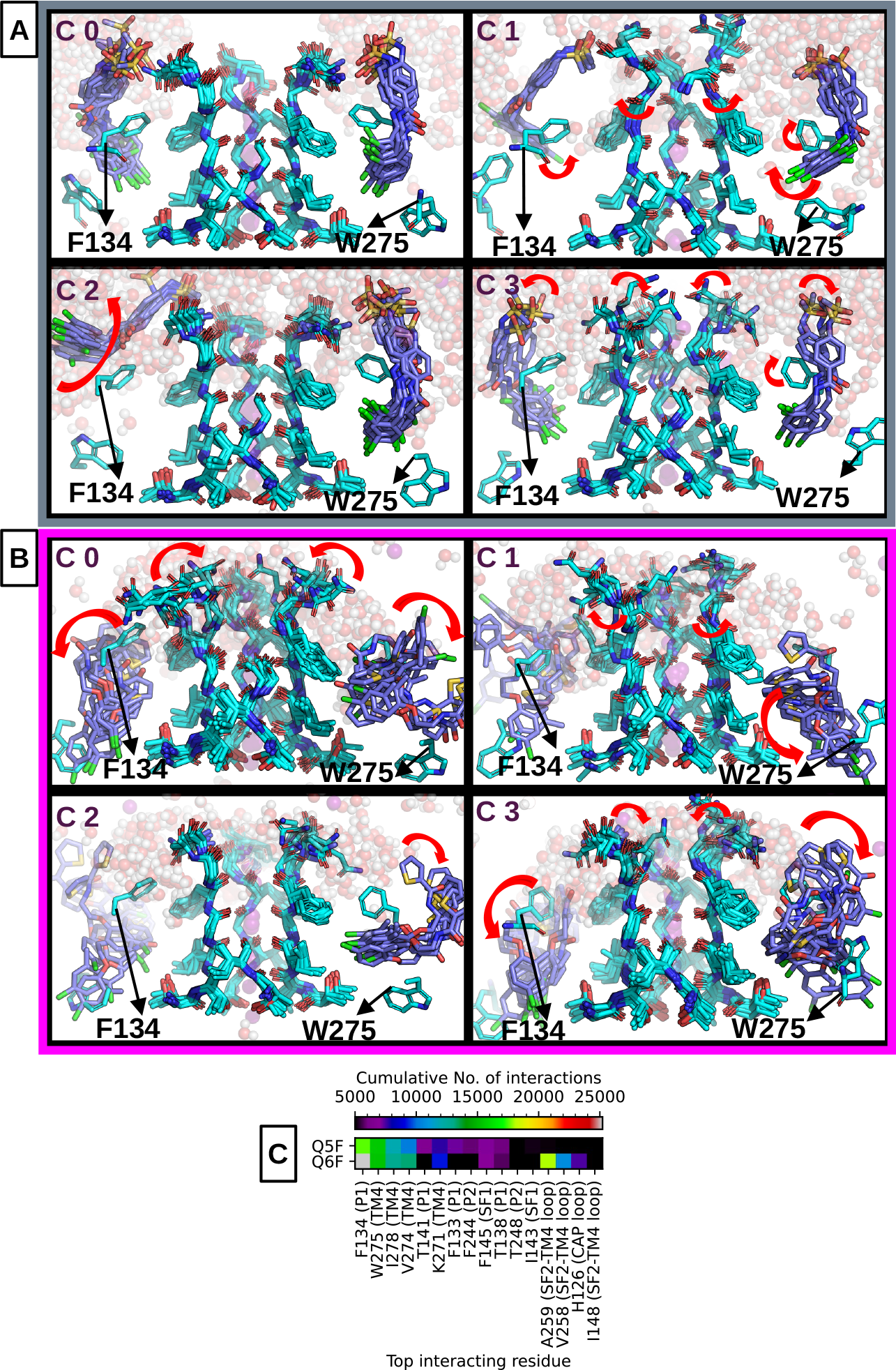
Representative snapshots of the cluster analysis carried out in Figure S4. SF2 residues 253 to 255 located in the foreground are not shown for clarity. This analysis is for Q6F **A**. and Q5F **B. C**. Most interacting residues in the cumulative interaction profile (N=10) with both ligands.

**Figure S6:**
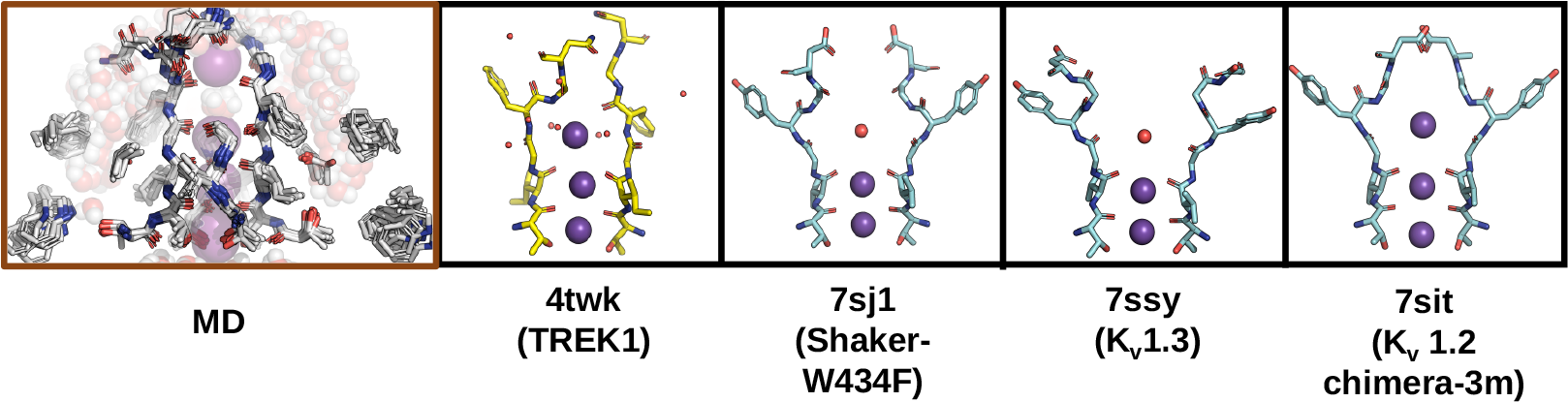
Comparison between a SF1 non-canonical conformation sampled by the apo-channel in our simulations, and structures of some K^+^ channels reported in the PDB showing a C-type inactivation.

**Figure S7:**
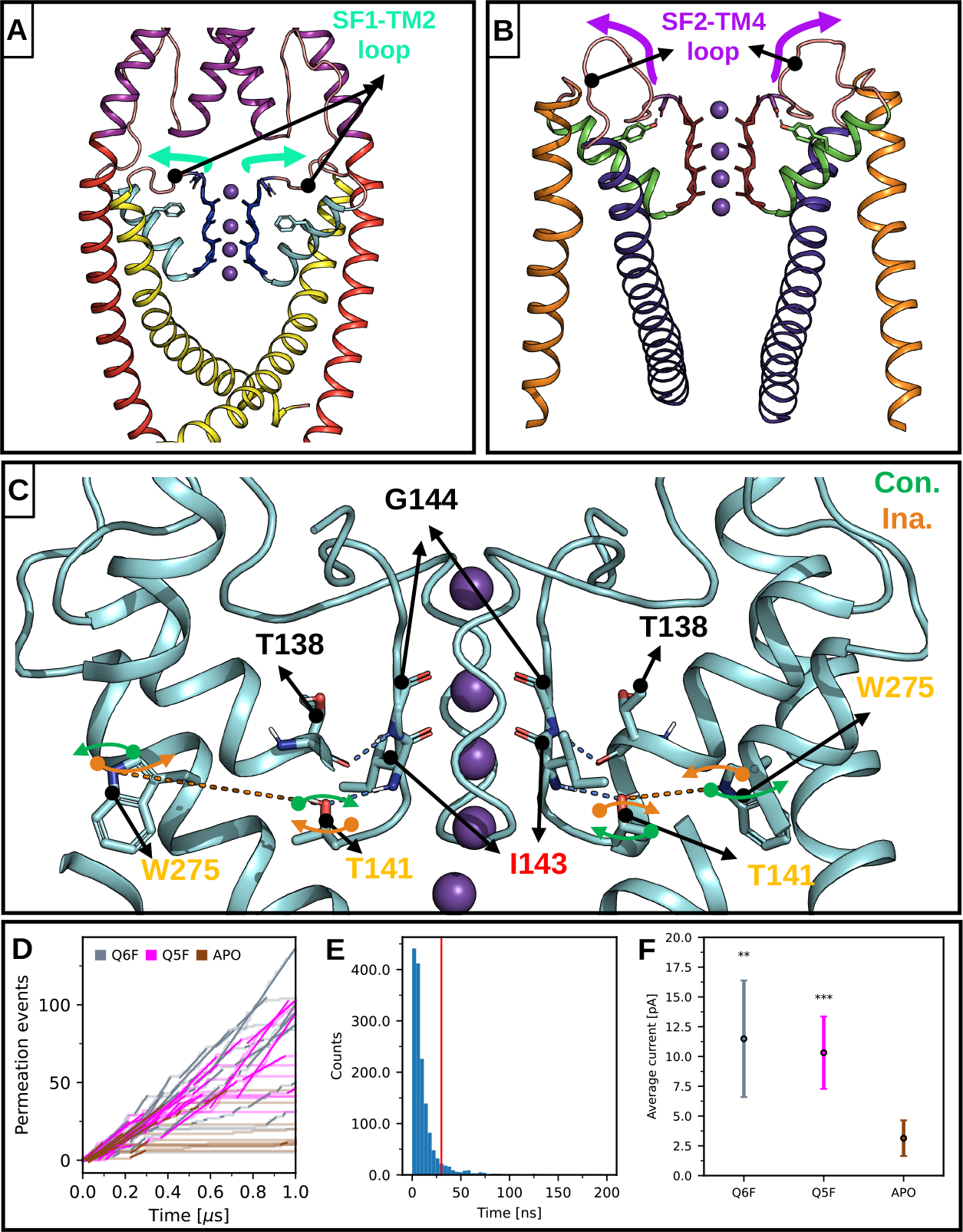
**A, B, C**. Three possible gating mechanisms converging on the SF according to Table 1. The color coding in the names corresponds to the same coding in Figure 2. **D**. Example of the calculation of the current in the channel when the filter is conducting: Multiple linear fits are made in the zones where the slope is different from zero, with the current when the filter is conducting being the average of these slopes. **E**. Distribution of time differences between consecutive permeation events in the total sampling. Taking a 30 ns cutoff, we cover approximately 90% of the total distribution; thus, only 10% of the permeations involve consecutive events taking more than 30 ns. **F**. Average channel current regardless of the conductive or non-conductive state of the SF (two-sided T-test for the means. ns: *P >* 0.05. *: *P ≤* 0.05. **: *P ≤* 0.01. ***: *P ≤* 0.001. Error bars correspond to confidence intervals of the mean).

**Figure S8:**
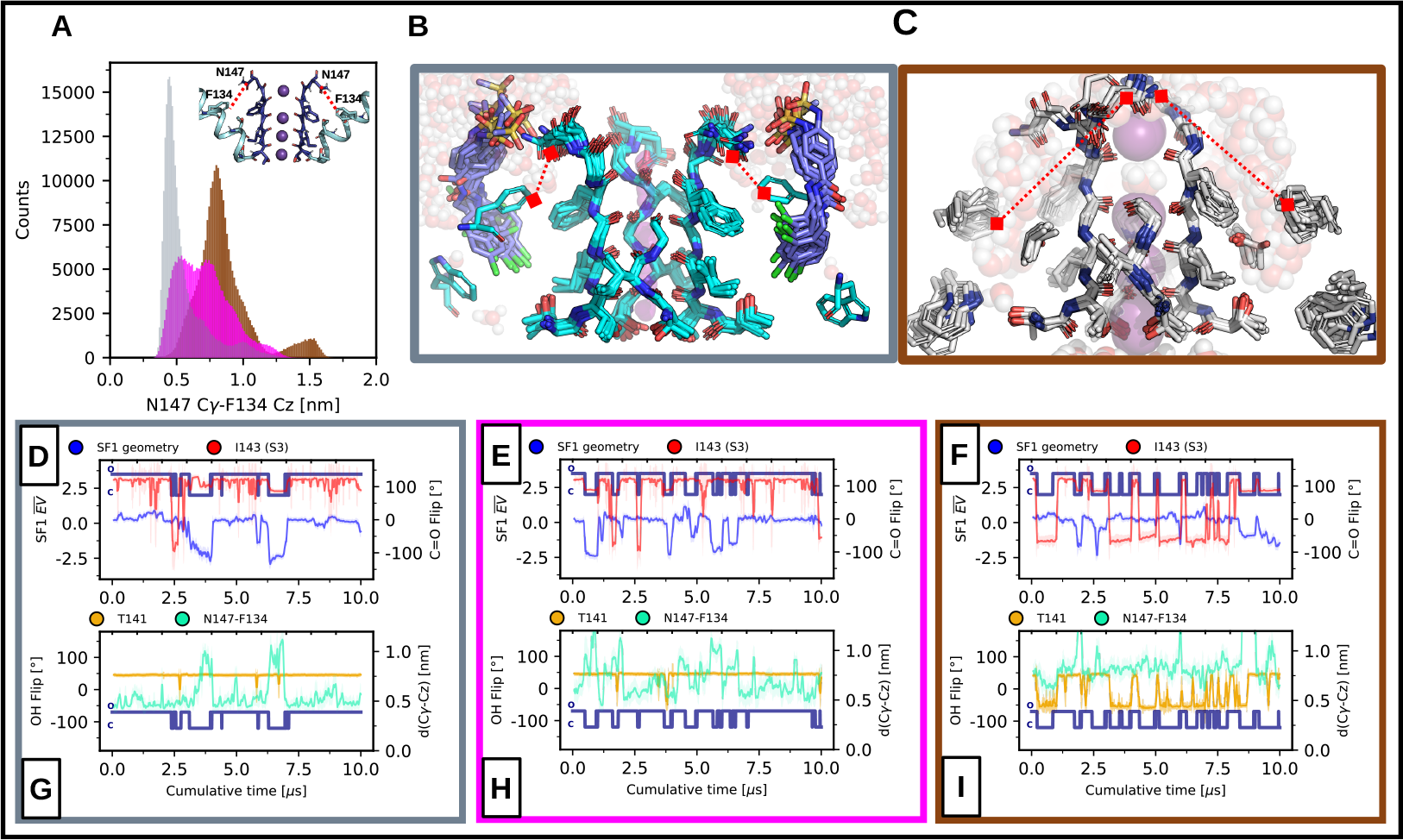
**A**. Distribution of distances between N147 (top of SF1) and F134 (P1 helix). These residues are equivalent to D80-W67 in KcsA, D447-W434 in *Shaker*, and D447-F434 in *Shaker* C-type inactivated mutant channels. The right panels show representative snapshots of the state of these distances in TREK1 in its Q6F-bound **B**. state and its apo **C**. state. **D, E, F**. Two gating mechanisms as a function of cumulative time (N=10): C-type inactivation distortions (given by the weighted average 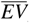); and the carbonyl flip in S3 (given by the N-Cα-C-O dihedral angle in residue I143). The blue-navy traces indicate when the channel is permeating (O) or not permeating (C) ions, respectively. Each panel corresponds to the two holo cases and the apo case, respectively. **G, H, I.)** Two order parameters about the modulation of TREK1 gating by ligands: Distance between N147 and F134 residues; and the stability of the HB network behind SF1 given by the χ1 angle of the T141 residue.

**Figure S9:**
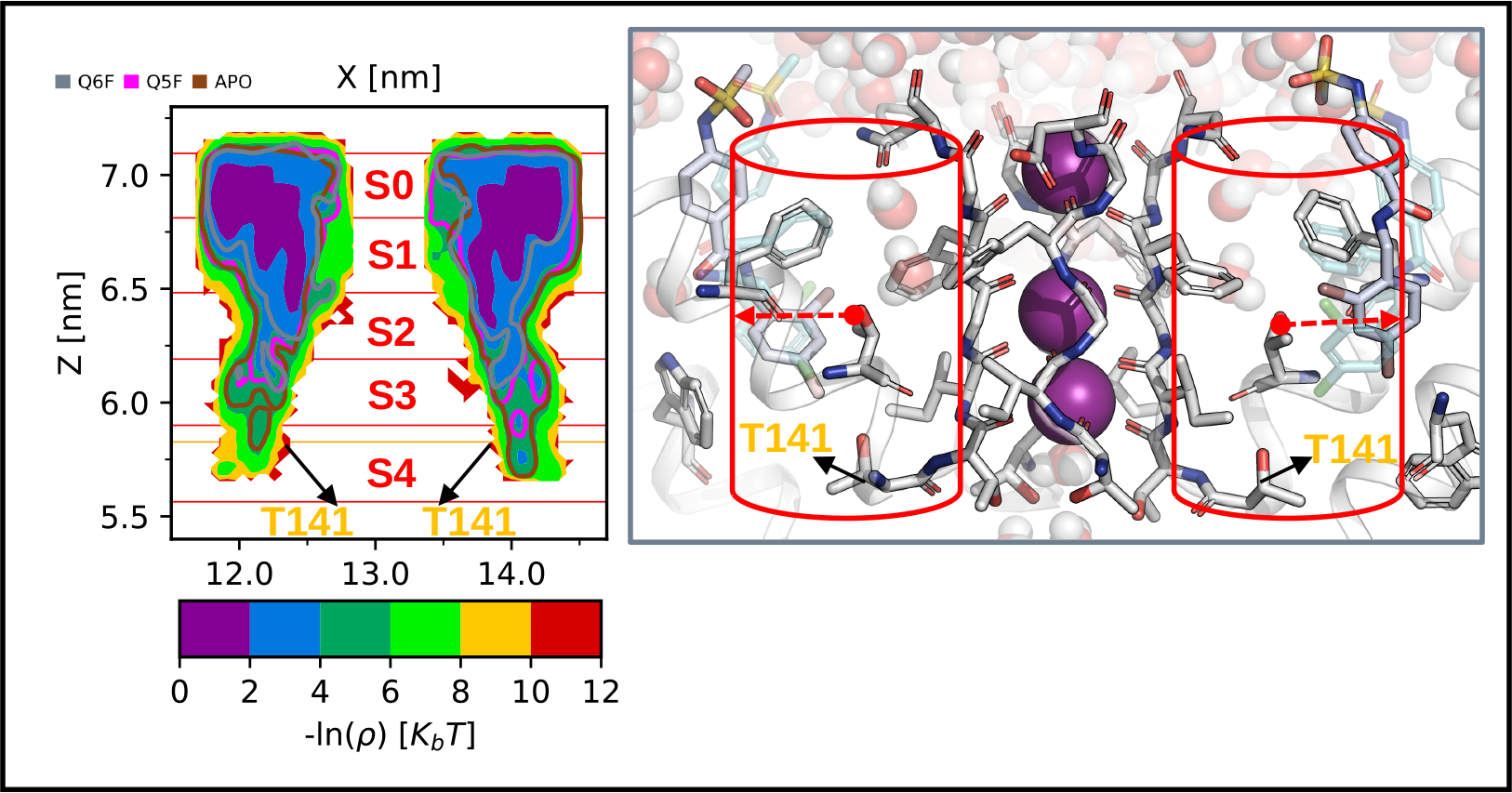
**Left panel:** Negative natural logarithm of the probability density function (PDF) of the oxygen positions of the water molecules in the total sample as a function of the X and Z axes. This density refers to water molecules within a cylinder of radius 0.5 nm behind SF1. The isocontours in the negative logarithm of the PDF indicate which regions are sampled by the channel depending on whether it is bound to ligands (Q6F, Q5F) or not (apo). The red lines indicate the positions of the carbonyls of the residues forming SF1, and the orange line denotes the position of the hydroxyl of residue T141 involved in the mostly ligand-regulated gating mechanism. **Right panel:** A representative snapshot of the limits used to obtain the probability density of water behind SF1.

**Figure S10:**
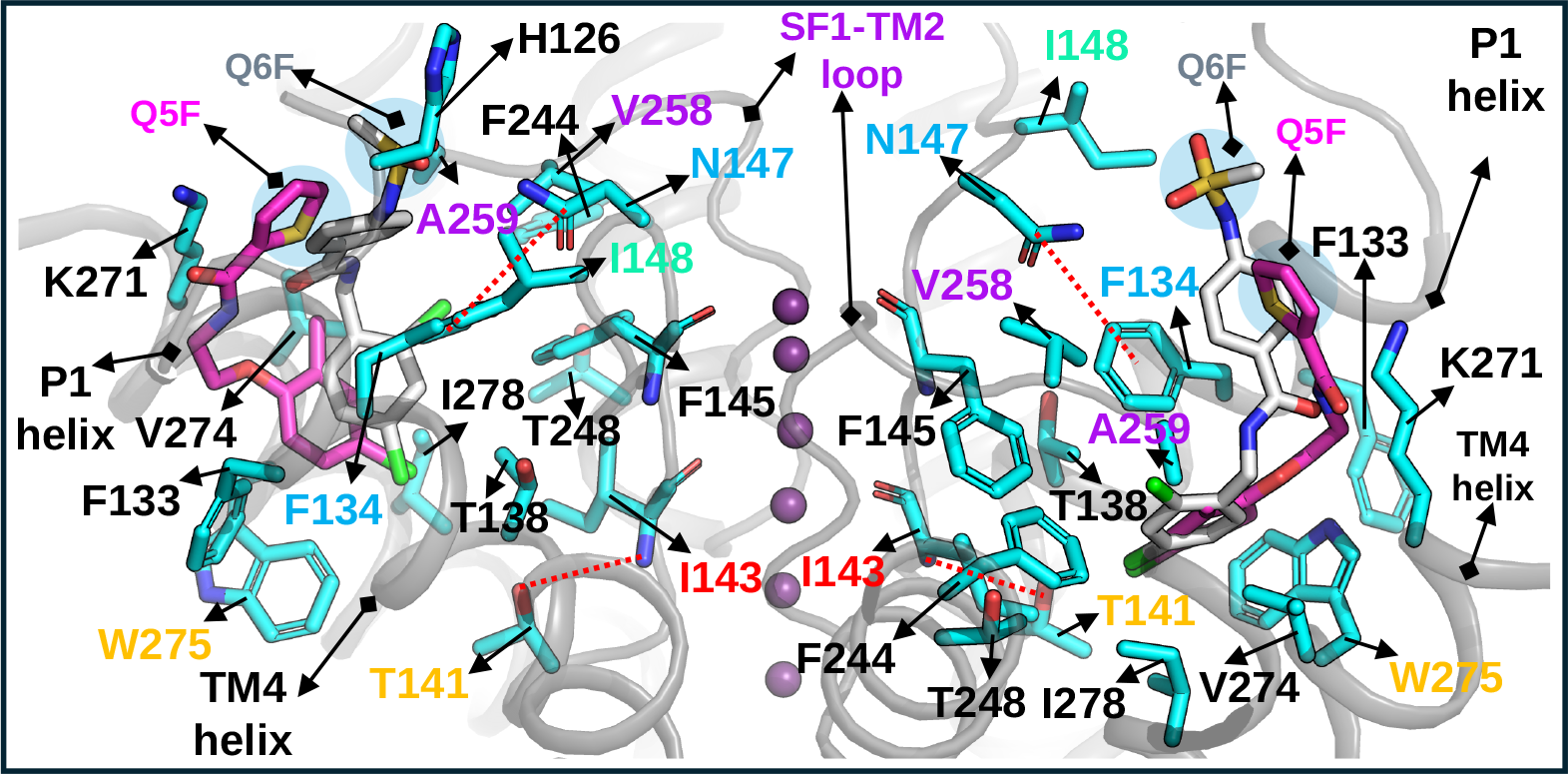
Representation of the residues that mostly interact with the Q6F and Q5F ligands as indicated in the table of Figure S6 C. Highlighted in red dotted lines are the interactions mostly affected by the ligands and that are involved in the two gating mechanisms: C-type inactivation and S3 flipping. It is observed that Q6F is longer than Q5F (due to its sulfonamide group) and interacts more strongly with N147 (which explains why Q6F is more effective in preventing C-type inactivations). Additionally, both ligands interact effectively with the lower part of SF1, which prevents the entry of water into this region (explaining why both ligands prevent non-permeation regimes due to S3 flipping).

**Figure S11:**
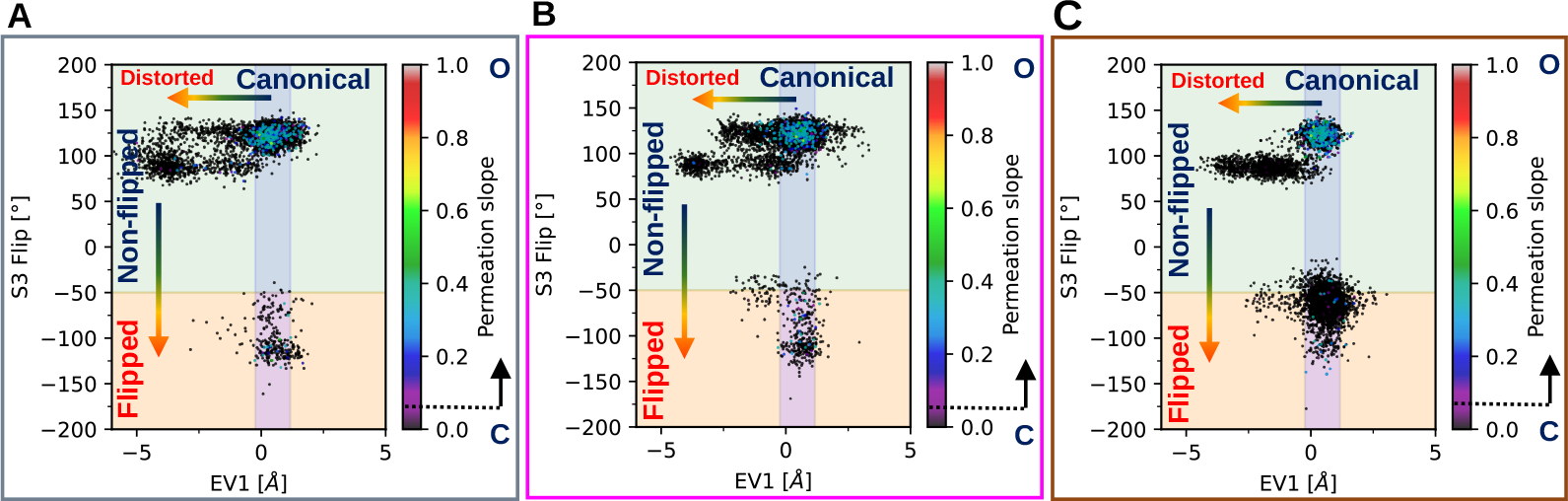
Coupling between two gating mechanisms: C-type inactivation and carbonyl flipping at S3 for the Q6F-bound **A**. Q5F-bound **B**. And apo channel **C**..

**Figure S12:**
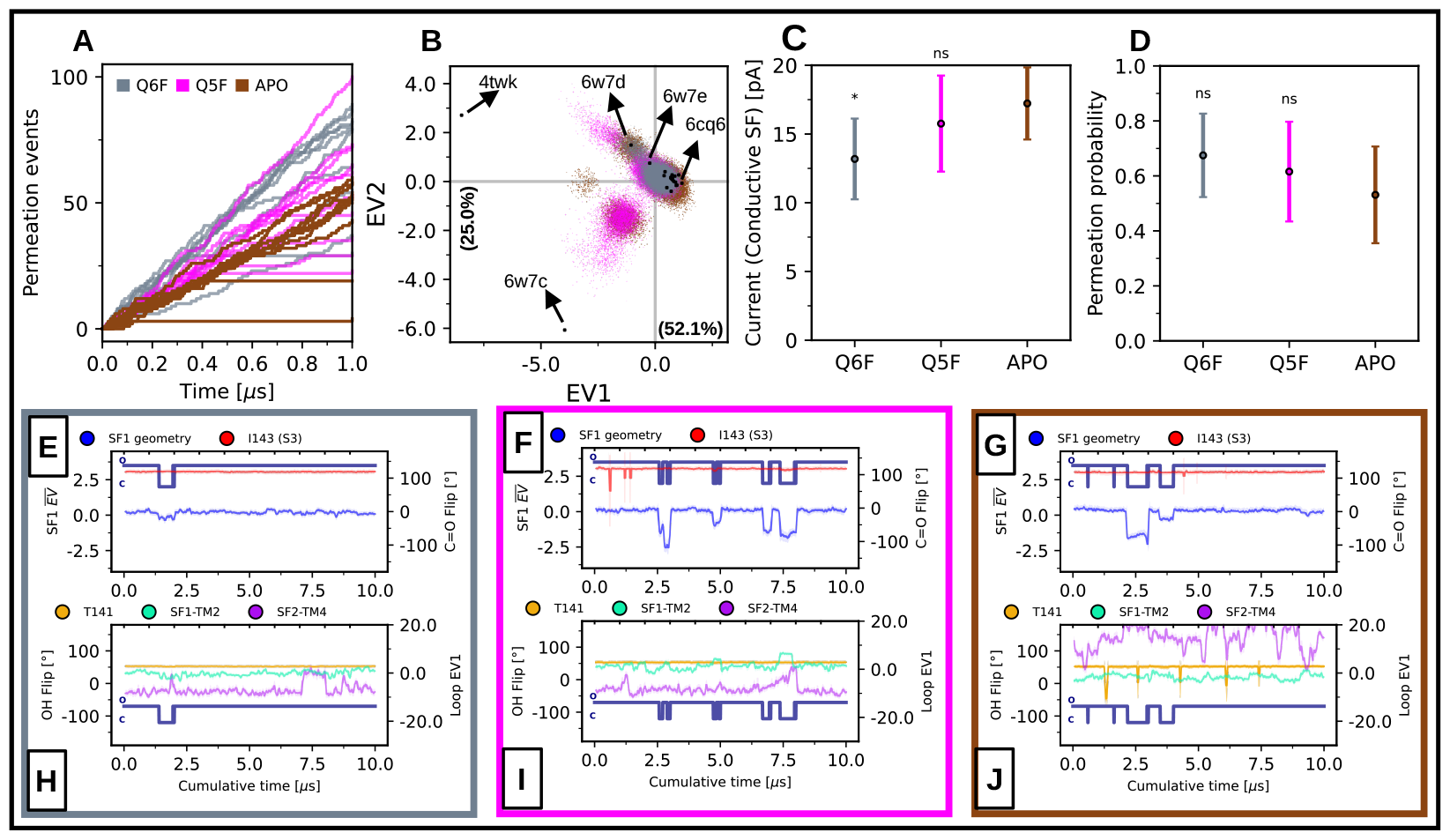
Results with the Amber force-field: **A**. Cumulative permeation events over 1 µs (N=10 for each condition). **B**. Total sampling of the holo and apo simulations projected on eigenvectors 1 and 2 of the SF1 main-chain experimental ensemble (30 µs of sampling are projected. EV means eigenvector). **D, C**. Permeation probability and channel current when the SF is conductive (See Methods). Both agree qualitatively with that observed in single trace currents experiments (two-sided T-test for the means. ns: P *>* 0.05. *: P *≤* 0.05. **: P *≤* 0.01. ***: P *≤* 0.001. Error bars correspond to confidence intervals of the mean). **E, F, G**. Two gating mechanisms as a function of cumulative time (N=10): C-type inactivation distortions (given by the weighted average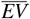); and the carbonyl flip in S3 (given by the N-Cα-C-O dihedral angle in residue I143). The blue-navy traces indicate when the channel is permeating (O) or not permeating (C) ions, respectively. Each panel corresponds to the two holo cases and the apo case, respectively. **H, I, J)** Three hypotheses about the modulation of TREK1 gating by ligands: Dynamics of the SF1-TM2 and SF2-TM4 loops given by the EV1 of their main-chain; and the stability of the HB network behind SF1 given by the χ1 angle of the T141 residue.

**Figure S13:**
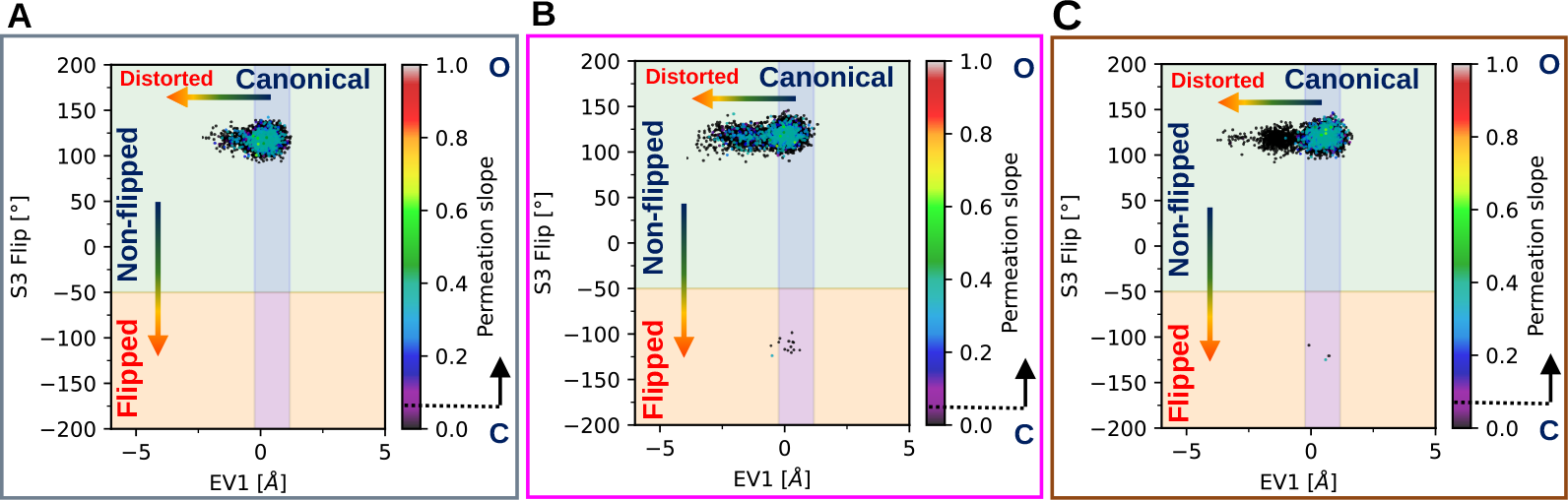
Coupling between two gating mechanisms using Amber force-field: C-type inactivation and carbonyl flipping at S3 for the Q6F-bound **A**. Q5F-bound **B**. And apo channel **C**..Table 1: Possible gating mechanisms converging in the SF.

## References

[1] Florian Lesage, Cécile Terrenoire, Georges Romey, and Michel Lazdunski. Human TREK2, a 2P domain mechano-sensitive K+ channel with multiple regulations by polyunsaturated fatty acids, lysophospholipids, and Gs, Gi, and Gq protein-coupled receptors. Journal of Biological Chemistry, 275(37):28398–28405, September 2000. ISSN 0021-9258. doi:10.1074/jbc.m002822200. URL http://dx.doi.org/10.1074/jbc.M002822200.

[2] Florian Lesage and Michel Lazdunski. Molecular and functional properties of two-pore-domain potassium channels. American Journal of Physiology-Renal Physiology, 279(5):F793–F801, November 2000. ISSN 1522-1466. doi:10.1152/ajprenal.2000.279.5.f793. URL http://dx.doi.org/10.1152/ajprenal.2000.279.5.F793.

[3] Steve AN Goldstein, Detlef Bockenhauer, Ita O’Kelly, and Noam Zilberberg. Potassium leak channels and the KCNK family of two-P-domain subunits. Nature Reviews Neuroscience, 2(3):175–184, March 2001. ISSN 1471-0048. doi:10.1038/35058574. URL http://dx.doi.org/10.1038/35058574.

[4] Vijay Renigunta, Günter Schlichthörl, and Jürgen Daut. Much more than a leak: structure and function of K2P-channels. Pflügers Archiv - European Journal of Physiology, 467(5):867–894, March 2015. ISSN 1432-2013. doi:10.1007/s00424-015-1703-7. URL http://dx.doi.org/10.1007/s00424-015-1703-7.

[5] Marcus Schewe, Ehsan Nematian-Ardestani, Han Sun, Marianne Musinszki, Sönke Cordeiro, Giovanna Bucci, Bert L de Groot, Stephen J Tucker, Markus Rapedius, and Thomas Baukrowitz. A non-canonical voltage-sensing mechanism controls gating in K2P K+ channels. Cell, 164(5):937–949, February 2016. ISSN 0092-8674. doi:10.1016/j.cell.2016.02.002. URL http://dx.doi.org/10.1016/j.cell.2016.02.002.

[6] Trevor A Richter, Gennady A Dvoryanchikov, Nirupa Chaudhari, and Stephen D Roper. Acid-sensitive two-pore domain potassium (K2P) channels in mouse taste buds. Journal of Neurophysiology, 92(3):1928–1936, September 2004. ISSN 1522-1598. doi:10.1152/jn.00273.2004. URL http://dx.doi.org/10.1152/jn.00273.2004.

[7] Dawon Kang, Changyong Choe, and Donghee Kim. Thermosensitivity of the two-pore domain K+ channels TREK-2 and TRAAK. The Journal of Physiology, 564(1):103–116, March 2005. ISSN 1469-7793. doi:10.1113/jphysiol.2004.081059. URL http://dx.doi.org/10.1113/jphysiol.2004.081059.

[8] Sviatoslav N Bagriantsev, Rémi Peyronnet, Kimberly A Clark, Eric Honoré, and Daniel L Minor Jr. Multiple modalities converge on a common gate to control K2P channel function. The EMBO Journal, 30(17):3594–3606, July 2011. ISSN 0261-4189. doi:10.1038/emboj.2011.230. URL http://dx.doi.org/10.1038/emboj.2011.230.

[9] KJ Buckler and E Honore. The lipid-activated two-pore domain K+ channel TREK-1 is resistant to hypoxia: implication for ischaemic neuroprotection. The Journal of Physiology, 562(1):213–222, December 2004. ISSN 1469-7793. doi:10.1113/jphysiol.2004.077503. URL http://dx.doi.org/10.1113/jphysiol.2004.077503.

[10] François Maingret, Amanda J Patel, Florian Lesage, Michel Lazdunski, and Eric Honoré. Lysophospholipids open the two-pore domain mechano-gated K+ channels TREK-1 and TRAAK. Journal of Biological Chemistry, 275 (14):10128–10133, April 2000. ISSN 0021-9258. doi:10.1074/jbc.275.14.10128. URL http://dx.doi.org/10.1074/jbc.275.14.10128.

[11] Jean Chemin, Amanda Jane Patel, Fabrice Duprat, Inger Lauritzen, Michel Lazdunski, and Eric Honoré. A phospholipid sensor controls mechanogating of the K+ channel TREK-1. The EMBO Journal, 24(1):44–53, December 2004. ISSN 1460-2075. doi:10.1038/sj.emboj.7600494. URL http://dx.doi.org/10.1038/sj.emboj.7600494.

[12] Eric Honoré, Amanda Jane Patel, Jean Chemin, Thomas Suchyna, and Frederick Sachs. Desensitization of mechano-gated K2P channels. Proceedings of the National Academy of Sciences, 103(18):6859–6864, May 2006. ISSN 1091-6490. doi:10.1073/pnas.0600463103. URL http://dx.doi.org/10.1073/pnas.0600463103.

[13] Joost HA Folgering, Reza Sharif-Naeini, Alexandra Dedman, Amanda Patel, Patrick Delmas, and Eric Honoré. Molecular basis of the mammalian pressure-sensitive ion channels: focus on vascular mechanotransduction. Progress in Biophysics and Molecular Biology, 97(2–3):180–195, June 2008. ISSN 0079-6107. doi:10.1016/j.pbiomolbio.2008.02.006. URL http://dx.doi.org/10.1016/j.pbiomolbio.2008.02.006.

[14] Prafulla Aryal, Viwan Jarerattanachat, Michael V Clausen, Marcus Schewe, Conor McClenaghan, Liam Argent, Linus J Conrad, Yin Y Dong, Ashley CW Pike, Elisabeth P Carpenter, et al. Bilayer-mediated structural transitions control mechanosensitivity of the trek-2 k2p channel. Structure, 25(5):708–718.e2, May 2017. ISSN 0969-2126. doi:10.1016/j.str.2017.03.006. URL http://dx.doi.org/10.1016/j.str.2017.03.006.

[15] Frederic Duprat, Christian Girard, G Jarretou, and M Lazdunski. Pancreatic two p domain K+ channels TALK-1 and TALK-2 are activated by nitric oxide and reactive oxygen species. The Journal of physiology, 562(1):235–244, December 2004. ISSN 1469-7793. doi:10.1113/jphysiol.2004.071266. URL http://dx.doi.org/10.1113/jphysiol.2004.071266.

[16] Eric Honoré. The neuronal background K2P channels: focus on TREK1. Nature Reviews Neuroscience, 8(4): 251–261, April 2007. ISSN 1471-0048. doi:10.1038/nrn2117. URL http://dx.doi.org/10.1038/nrn2117.

[17] Alaeddine Djillani, Jean Mazella, Catherine Heurteaux, and Marc Borsotto. Role of TREK-1 in health and disease, focus on the central nervous system. Frontiers in Pharmacology, 10, April 2019. ISSN 1663-9812. doi:10.3389/fphar.2019.00379. URL http://dx.doi.org/10.3389/fphar.2019.00379.

[18] Lianne Pope and Daniel L Minor Jr. The Polysite Pharmacology of TREK K2P Channels, page 51–65. Springer Nature Singapore, 2021. ISBN 9789811642548. doi:10.1007/978-981-16-4254-8_4. URL http://dx.doi.org/10.1007/978-981-16-4254-8_4.

[19] Andrew M Natale, Parker E Deal, and Daniel L Minor Jr. Structural insights into the mechanisms and pharmacology of K2P potassium channels. Journal of Molecular Biology, 433(17):166995, August 2021. ISSN 0022-2836. doi:10.1016/j.jmb.2021.166995. URL http://dx.doi.org/10.1016/j.jmb.2021.166995.

[20] Alistair Mathie. Ion channels as novel therapeutic targets in the treatment of pain. Journal of Pharmacy and Pharmacology, 62(9):1089–1095, August 2010. ISSN 2042-7158. doi:10.1111/j.2042-7158.2010.01131.x. URL http://dx.doi.org/10.1111/j.2042-7158.2010.01131.x.

[21] Marco Lolicato, Andrew M Natale, Fayal Abderemane-Ali, David Crottès, Sara Capponi, Ramona Duman, Armin Wagner, John M Rosenberg, Michael Grabe, and Daniel L Minor Jr. K2P channel C-type gating involves asymmetric selectivity filter order-disorder transitions. Science Advances, 6(44), October 2020. ISSN 2375-2548. doi:10.1126/sciadv.abc9174. URL http://dx.doi.org/10.1126/sciadv.abc9174.

[22] Marco Lolicato, Cristina Arrigoni, Takahiro Mori, Yoko Sekioka, Clifford Bryant, Kimberly A Clark, and Daniel L Minor Jr. K2P2.1 (TREK-1)–activator complexes reveal a cryptic selectivity filter binding site. Nature, 547 (7663):364–368, July 2017. ISSN 1476-4687. doi:10.1038/nature22988. URL http://dx.doi.org/10.1038/nature22988.

[23] Lianne Pope, Marco Lolicato, and Daniel L Minor. Polynuclear ruthenium amines inhibit K2P channels via a “Finger in the Dam” mechanism. Cell Chemical Biology, 27(5):511–524.e4, May 2020. ISSN 2451-9456. doi:10.1016/j.chembiol.2020.01.011. URL http://dx.doi.org/10.1016/j.chembiol.2020.01.011.

[24] Marcus Schewe, Han Sun, Ümit Mert, Alexandra Mackenzie, Ashley CW Pike, Friederike Schulz, Cristina Constantin, Kirsty S Vowinkel, Linus J Conrad, Aytug K Kiper, et al. A pharmacological master key mechanism that unlocks the selectivity filter gate in K+ channels. Science, 363(6429):875–880, February 2019. ISSN 1095-9203. doi:10.1126/science.aav0569. URL http://dx.doi.org/10.1126/science.aav0569.

[25] Philipp AM Schmidpeter, John T Petroff, Leila Khajoueinejad, Aboubacar Wague, Cheryl Frankfater, Wayland WL Cheng, Crina M Nimigean, and Paul M Riegelhaupt. Membrane phospholipids control gating of the mechanosensitive potassium leak channel TREK1. Nature Communications, 14(1), February 2023. ISSN 2041-1723. doi:10.1038/s41467-023-36765-w. URL http://dx.doi.org/10.1038/s41467-023-36765-w.

[26] Michel Fink, Fabrice Duprat, Florian Lesage, Roberto Reyes, Georges Romey, Catherine Heurteaux, and M Lazdunski. Cloning, functional expression and brain localization of a novel unconventional outward rectifier K+ channel. The EMBO journal, 15(24):6854–6862, 1996. doi:10.1002/j.1460-2075.1996.tb01077.x. URL https://www.embopress.org/doi/abs/10.1002/j.1460-2075.1996.tb01077.x.

[27] Marco Lolicato, Paul M Riegelhaupt, Cristina Arrigoni, Kimberly A Clark, and Daniel L Minor. Transmembrane helix straightening and buckling underlies activation of mechanosensitive and thermosensitive K2P channels. Neuron, 84(6):1198–1212, December 2014. ISSN 0896-6273. doi:10.1016/j.neuron.2014.11.017. URL http://dx.doi.org/10.1016/j.neuron.2014.11.017.

[28] Julian T Brennecke and Bert L de Groot. Mechanism of mechanosensitive gating of the TREK-2 potassium channel. Biophysical Journal, 114(6):1336–1343, March 2018. ISSN 0006-3495. doi:10.1016/j.bpj.2018.01.030. URL http://dx.doi.org/10.1016/j.bpj.2018.01.030.

[29] Peter Proks, Marcus Schewe, Linus J Conrad, Shanlin Rao, Kristin Rathje, Karin EJ Rödström, Elisabeth P Carpenter, Thomas Baukrowitz, and Stephen J Tucker. Norfluoxetine inhibits TREK-2 K2P channels by multiple mechanisms including state-independent effects on the selectivity filter gate. Journal of General Physiology, 153(8), May 2021. ISSN 1540-7748. doi:10.1085/jgp.202012812. URL http://dx.doi.org/10.1085/jgp.202012812.

[30] Conor McClenaghan, Marcus Schewe, Prafulla Aryal, Elisabeth P Carpenter, Thomas Baukrowitz, and Stephen J Tucker. Polymodal activation of the TREK-2 K2P channel produces structurally distinct open states. Journal of General Physiology, 147(6):497–505, May 2016. ISSN 1540-7748. doi:10.1085/jgp.201611601. URL http://dx.doi.org/10.1085/jgp.201611601.

[31] Yin Yao Dong, Ashley CW Pike, Alexandra Mackenzie, Conor McClenaghan, Prafulla Aryal, Liang Dong, Andrew Quigley, Mariana Grieben, Solenne Goubin, Shubhashish Mukhopadhyay, et al. K2P channel gating mechanisms revealed by structures of TREK-2 and a complex with prozac. Science, 347(6227):1256–1259, March 2015. ISSN 1095-9203. doi:10.1126/science.1261512. URL http://dx.doi.org/10.1126/science.1261512.

[32] Robert A Rietmeijer, Ben Sorum, Baobin Li, and Stephen G Brohawn. Physical basis for distinct basal and mechanically gated activity of the human K+ channel TRAAK. Neuron, 109(18):2902–2913.e4, September 2021. ISSN 0896-6273. doi:10.1016/j.neuron.2021.07.009. URL http://dx.doi.org/10.1016/j.neuron.2021.07.009.

[33] Paula L Piechotta, Markus Rapedius, Phillip J Stansfeld, Murali K Bollepalli, Gunter Erhlich, Isabelle Andres-Enguix, Hariolf Fritzenschaft, Niels Decher, Mark SP Sansom, Stephen J Tucker, and Thomas Baukrowitz. The pore structure and gating mechanism of K2P channels. The EMBO Journal, 30(17):3607–3619, August 2011. ISSN 0261-4189. doi:10.1038/emboj.2011.268. URL http://dx.doi.org/10.1038/emboj.2011.268.

[34] Markus Rapedius, Matthias R Schmidt, Chetan Sharma, Phillip J Stansfeld, Mark SP Sansom, Thomas Baukrowitz, and Stephen J Tucker. State-independent intracellular access of quaternary ammonium blockers to the pore of TREK-1. Channels, 6(6):473–478, November 2012. ISSN 1933-6969. doi:10.4161/chan.22153. URL http://dx.doi.org/10.4161/chan.22153.

[35] Ismail Ben Soussia, Frank S Choveau, Sandy Blin, Eun-Jin Kim, Sylvain Feliciangeli, Franck C Chatelain, Dawon Kang, Delphine Bichet, and Florian Lesage. Antagonistic effect of a cytoplasmic domain on the basal activity of polymodal potassium channels. Frontiers in Molecular Neuroscience, 11, September 2018. ISSN 1662-5099. doi:10.3389/fnmol.2018.00301. URL http://dx.doi.org/10.3389/fnmol.2018.00301.

[36] Frank S Choveau, Ismail Ben Soussia, Delphine Bichet, Chatelain C Franck, Sylvain Feliciangeli, and Florian Lesage. Convergence of multiple stimuli to a single gate in TREK1 and TRAAK potassium channels. Frontiers in Pharmacology, 12, September 2021. ISSN 1663-9812. doi:10.3389/fphar.2021.755826. URL http://dx.doi.org/10.3389/fphar.2021.755826.

[37] Noam Zilberberg, Nitza Ilan, and Steve AN Goldstein. Kcnkø: opening and closing the 2-p-domain potassium leak channel entails “C-type” gating of the outer pore. Neuron, 32(4):635–648, November 2001. ISSN 0896-6273. doi:10.1016/s0896-6273(01)00503-7. URL http://dx.doi.org/10.1016/S0896-6273(01)00503-7.

[38] Asi Cohen, Yuval Ben-Abu, Shelly Hen, and Noam Zilberberg. A novel mechanism for human K2P2.1 channel gating: facilitation of C-type gating by protonation of extracellular histidine residues. Journal of Biological Chemistry, 283(28):19448–19455, July 2008. ISSN 0021-9258. doi:10.1074/jbc.m801273200. URL http://dx.doi.org/10.1074/jbc.M801273200.

[39] Qiansen Zhang, Jie Fu, Shaoying Zhang, Peipei Guo, Shijie Liu, Juwen Shen, Jiangtao Guo, Huaiyu Yang, and Xuebiao Yao. ‘C-type’ closed state and gating mechanisms of K2P channels revealed by conformational changes of the TREK-1 channel. Journal of Molecular Cell Biology, 14(1), January 2022. ISSN 1759-4685. doi:10.1093/jmcb/mjac002. URL http://dx.doi.org/10.1093/jmcb/mjac002.

[40] Jumin Lee, Xi Cheng, Jason M. Swails, Min Sun Yeom, Peter K. Eastman, Justin A. Lemkul, Shuai Wei, Joshua Buckner, Jong Cheol Jeong, Yifei Qi, Sunhwan Jo, Vijay S. Pande, David A. Case, Charles L. Brooks, Alexander D. MacKerell, Jeffery B. Klauda, and Wonpil Im. Charmm-gui input generator for namd, gromacs, amber, openmm, and charmm/openmm simulations using the charmm36 additive force field. Journal of Chemical Theory and Computation, 12(1):405–413, December 2015. ISSN 1549-9626. doi:10.1021/acs.jctc.5b00935. URL http://dx.doi.org/10.1021/acs.jctc.5b00935.

[41] Sunhwan Jo, Taehoon Kim, Vidyashankara G. Iyer, and Wonpil Im. Charmm-gui: A web-based graphical user interface for charmm. Journal of Computational Chemistry, 29(11):1859–1865, June 2008. ISSN 1096-987X. doi:10.1002/jcc.20945. URL http://dx.doi.org/10.1002/jcc.20945.

[42] Sunhwan Jo, Joseph B. Lim, Jeffery B. Klauda, and Wonpil Im. Charmm-gui membrane builder for mixed bilayers and its application to yeast membranes. Biophysical Journal, 97(1):50–58, July 2009. ISSN 0006-3495. doi:10.1016/j.bpj.2009.04.013. URL http://dx.doi.org/10.1016/j.bpj.2009.04.013.

[43] Emilia L. Wu, Xi Cheng, Sunhwan Jo, Huan Rui, Kevin C. Song, Eder M. Dávila-Contreras, Yifei Qi, Jumin Lee, Viviana Monje-Galvan, Richard M. Venable, Jeffery B. Klauda, and Wonpil Im. Charmm-guimembrane buildertoward realistic biological membrane simulations. Journal of Computational Chemistry, 35(27):1997–2004, August 2014. ISSN 0192-8651. doi:10.1002/jcc.23702. URL http://dx.doi.org/10.1002/jcc.23702.

[44] Herman JC Berendsen, David van der Spoel, and Rudi van Drunen. Gromacs: A message-passing parallel molecular dynamics implementation. Computer Physics Communications, 91(1–3):43–56, September 1995. ISSN 0010-4655. doi:10.1016/0010-4655(95)00042-e. URL http://dx.doi.org/10.1016/0010-4655(95)00042-E.

[45] Erik Lindahl, Berk Hess, and David Van Der Spoel. Gromacs 3.0: a package for molecular simulation and trajectory analysis. Journal of Molecular Modeling, 7(8):306–317, August 2001. ISSN 0948-5023. doi:10.1007/s008940100045. URL http://dx.doi.org/10.1007/s008940100045.

[46] David Van Der Spoel, Erik Lindahl, Berk Hess, Gerrit Groenhof, Alan E Mark, and Herman JC Berendsen. Gromacs: fast, flexible, and free. Journal of Computational Chemistry, 26(16):1701–1718, October 2005. ISSN 1096-987X. doi:10.1002/jcc.20291. URL http://dx.doi.org/10.1002/jcc.20291.

[47] Berk Hess, Carsten Kutzner, David van der Spoel, and Erik Lindahl. Gromacs 4: Algorithms for highly efficient, load-balanced, and scalable molecular simulation. Journal of Chemical Theory and Computation, 4(3):435–447, February 2008. ISSN 1549-9626. doi:10.1021/ct700301q. URL http://dx.doi.org/10.1021/ct700301q.

[48] Sander Pronk, Szilárd Páll, Roland Schulz, Per Larsson, Pär Bjelkmar, Rossen Apostolov, Michael R. Shirts, Jeremy C. Smith, Peter M. Kasson, David van der Spoel, Berk Hess, and Erik Lindahl. Gromacs 4.5: a high-throughput and highly parallel open source molecular simulation toolkit. Bioinformatics, 29(7):845–854, February 2013. ISSN 1367-4803. doi:10.1093/bioinformatics/btt055. URL http://dx.doi.org/10.1093/bioinformatics/btt055.

[49] Mark James Abraham, Teemu Murtola, Roland Schulz, Szilárd Páll, Jeremy C. Smith, Berk Hess, and Erik Lindahl. Gromacs: High performance molecular simulations through multi-level parallelism from laptops to supercomputers. SoftwareX, 1–2:19–25, September 2015. ISSN 2352-7110. doi:10.1016/j.softx.2015.06.001. URL http://dx.doi.org/10.1016/j.softx.2015.06.001.

[50] Jing Huang, Sarah Rauscher, Grzegorz Nawrocki, Ting Ran, Michael Feig, Bert L de Groot, Helmut Grubmüller, and Alexander D MacKerell. Charmm36m: an improved force field for folded and intrinsically disordered proteins. Nature Methods, 14(1):71–73, November 2016. ISSN 1548-7105. doi:10.1038/nmeth.4067. URL http://dx.doi.org/10.1038/nmeth.4067.

[51] A. D. MacKerell, D. Bashford, M. Bellott, R. L. Dunbrack, J. D. Evanseck, M. J. Field, S. Fischer, J. Gao, H. Guo, S. Ha, D. Joseph-McCarthy, L. Kuchnir, K. Kuczera, F. T. K. Lau, C. Mattos, S. Michnick, T. Ngo, D. T. Nguyen, B. Prodhom, W. E. Reiher, B. Roux, M. Schlenkrich, J. C. Smith, R. Stote, J. Straub, M. Watanabe, J. Wiórkiewicz-Kuczera, D. Yin, and M. Karplus. All-atom empirical potential for molecular modeling and dynamics studies of proteins. The Journal of Physical Chemistry B, 102(18):3586–3616, April 1998. ISSN 1520-5207. doi:10.1021/jp973084f. URL http://dx.doi.org/10.1021/jp973084f.

[52] Jeffery B. Klauda, Viviana Monje, Taehoon Kim, and Wonpil Im. Improving the charmm force field for polyunsaturated fatty acid chains. The Journal of Physical Chemistry B, 116(31):9424–9431, July 2012. ISSN 1520-5207. doi:10.1021/jp304056p. URL http://dx.doi.org/10.1021/jp304056p.

[53] Jeffery B. Klauda, Richard M. Venable, J. Alfredo Freites, Joseph W. O’Connor, Douglas J. Tobias, Carlos Mondragon-Ramirez, Igor Vorobyov, Alexander D. MacKerell, and Richard W. Pastor. Update of the charmm all-atom additive force field for lipids: Validation on six lipid types. The Journal of Physical Chemistry B, 114 (23):7830–7843, May 2010. ISSN 1520-5207. doi:10.1021/jp101759q. URL http://dx.doi.org/10.1021/jp101759q.

[54] K. Vanommeslaeghe, E. Hatcher, C. Acharya, S. Kundu, S. Zhong, J. Shim, E. Darian, O. Guvench, P. Lopes, I. Vorobyov, and A. D. Mackerell. Charmm general force field: A force field for drug-like molecules compatible with the charmm all-atom additive biological force fields. Journal of Computational Chemistry, 31(4):671–690, July 2009. ISSN 1096-987X. doi:10.1002/jcc.21367. URL http://dx.doi.org/10.1002/jcc.21367.

[55] K. Vanommeslaeghe and A. D. MacKerell. Automation of the charmm general force field (cgenff) i: Bond perception and atom typing. Journal of Chemical Information and Modeling, 52(12):3144–3154, November 2012. ISSN 1549-960X. doi:10.1021/ci300363c. URL http://dx.doi.org/10.1021/ci300363c.

[56] K. Vanommeslaeghe, E. Prabhu Raman, and A. D. MacKerell. Automation of the charmm general force field (cgenff) ii: Assignment of bonded parameters and partial atomic charges. Journal of Chemical Information and Modeling, 52(12):3155–3168, November 2012. ISSN 1549-960X. doi:10.1021/ci3003649. URL http://dx.doi.org/10.1021/ci3003649.

[57] Wenbo Yu, Xibing He, Kenno Vanommeslaeghe, and Alexander D. MacKerell. Extension of the charmm general force field to sulfonyl-containing compounds and its utility in biomolecular simulations. Journal of Computational Chemistry, 33(31):2451–2468, July 2012. ISSN 1096-987X. doi:10.1002/jcc.23067. URL http://dx.doi.org/10.1002/jcc.23067.

[58] Dmitrii Beglov and Benoît Roux. Finite representation of an infinite bulk system: Solvent boundary potential for computer simulations. The Journal of Chemical Physics, 100(12):9050–9063, June 1994. ISSN 1089-7690. doi:10.1063/1.466711. URL http://dx.doi.org/10.1063/1.466711.

[59] James A. Maier, Carmenza Martinez, Koushik Kasavajhala, Lauren Wickstrom, Kevin E. Hauser, and Carlos Simmerling. ff14sb: Improving the accuracy of protein side chain and backbone parameters from ff99sb. Journal of Chemical Theory and Computation, 11(8):3696–3713, July 2015. ISSN 1549-9626. doi:10.1021/acs.jctc.5b00255. URL http://dx.doi.org/10.1021/acs.jctc.5b00255.

[60] William L. Jorgensen, Jayaraman Chandrasekhar, Jeffry D. Madura, Roger W. Impey, and Michael L. Klein. Comparison of simple potential functions for simulating liquid water. The Journal of Chemical Physics, 79(2): 926–935, July 1983. ISSN 1089-7690. doi:10.1063/1.445869. URL http://dx.doi.org/10.1063/1.445869.

[61] Joakim P. M. Jämbeck and Alexander P. Lyubartsev. Derivation and systematic validation of a refined all-atom force field for phosphatidylcholine lipids. The Journal of Physical Chemistry B, 116(10):3164–3179, March 2012. ISSN 1520-5207. doi:10.1021/jp212503e. URL http://dx.doi.org/10.1021/jp212503e.

[62] Joakim P. M. Jämbeck and Alexander P. Lyubartsev. An extension and further validation of an all-atomistic force field for biological membranes. Journal of Chemical Theory and Computation, 8(8):2938–2948, July 2012. ISSN 1549-9626. doi:10.1021/ct300342n. URL http://dx.doi.org/10.1021/ct300342n.

[63] Junmei Wang, Wei Wang, Peter A. Kollman, and David A. Case. Automatic atom type and bond type perception in molecular mechanical calculations. Journal of Molecular Graphics and Modelling, 25(2):247–260, October 2006. ISSN 1093-3263. doi:10.1016/j.jmgm.2005.12.005. URL http://dx.doi.org/10.1016/j.jmgm.2005.12.005.

[64] Junmei Wang, Romain M. Wolf, James W. Caldwell, Peter A. Kollman, and David A. Case. Development and testing of a general amber force field. Journal of Computational Chemistry, 25(9):1157–1174, April 2004. ISSN 1096-987X. doi:10.1002/jcc.20035. URL http://dx.doi.org/10.1002/jcc.20035.

[65] Xibing He, Viet H. Man, Wei Yang, Tai-Sung Lee, and Junmei Wang. A fast and high-quality charge model for the next generation general amber force field. The Journal of Chemical Physics, 153(11), September 2020. ISSN 1089-7690. doi:10.1063/5.0019056. URL http://dx.doi.org/10.1063/5.0019056.

[66] In Suk Joung and Thomas E. Cheatham. Determination of alkali and halide monovalent ion parameters for use in explicitly solvated biomolecular simulations. The Journal of Physical Chemistry B, 112(30):9020–9041, July 2008. ISSN 1520-5207. doi:10.1021/jp8001614. URL http://dx.doi.org/10.1021/jp8001614.

[67] Alan W Sousa da Silva and Wim F Vranken. Acpype - antechamber python parser interface. BMC Research Notes, 5(1), July 2012. ISSN 1756-0500. doi:10.1186/1756-0500-5-367. URL http://dx.doi.org/10.1186/1756-0500-5-367.

[68] William G. Hoover. Canonical dynamics: Equilibrium phase-space distributions. Physical Review A, 31(3): 1695–1697, March 1985. ISSN 0556-2791. doi:10.1103/physreva.31.1695. URL http://dx.doi.org/10.1103/PhysRevA.31.1695.

[69] M. Parrinello and A. Rahman. Polymorphic transitions in single crystals: A new molecular dynamics method. Journal of Applied Physics, 52(12):7182–7190, December 1981. ISSN 1089-7550. doi:10.1063/1.328693. URL http://dx.doi.org/10.1063/1.328693.

[70] Berk Hess, Henk Bekker, Herman J. C. Berendsen, and Johannes G. E. M. Fraaije. Lincs: A linear constraint solver for molecular simulations. Journal of Computational Chemistry, 18(12):1463–1472, September 1997. ISSN 1096-987X. doi:10.1002/(sici)1096-987x(199709)18:12<1463::aid-jcc4>3.0.co;2-h. URL http://dx.doi.org/10.1002/(SICI)1096-987X(199709)18:12<1463::AID-JCC4>3.0.CO;2-H.

[71] Tom Darden, Darrin York, and Lee Pedersen. Particle mesh ewald: An N*log(N) method for ewald sums in large systems. The Journal of Chemical Physics, 98(12):10089–10092, June 1993. ISSN 1089-7690. doi:10.1063/1.464397. URL http://dx.doi.org/10.1063/1.464397.

[72] Benoît Roux. The membrane potential and its representation by a constant electric field in computer simulations. Biophysical Journal, 95(9):4205–4216, November 2008. ISSN 0006-3495. doi:10.1529/biophysj.108.136499. URL http://dx.doi.org/10.1529/biophysj.108.136499.

[73] James Gumbart, Fatemeh Khalili-Araghi, Marcos Sotomayor, and Benoît Roux. Constant electric field simulations of the membrane potential illustrated with simple systems. Biochimica et Biophysica Acta (BBA) - Biomembranes, 1818(2):294–302, February 2012. ISSN 0005-2736. doi:10.1016/j.bbamem.2011.09.030. URL http://dx.doi.org/10.1016/j.bbamem.2011.09.030.

[74] Travis E Oliphant. Python for scientific computing. Computing in science & engineering, 9(3):10–20, 2007.

[75] Charles R. Harris, K. Jarrod Millman, Stéfan J. van der Walt, Ralf Gommers, Pauli Virtanen, David Cournapeau, Eric Wieser, Julian Taylor, Sebastian Berg, Nathaniel J. Smith, Robert Kern, Matti Picus, Stephan Hoyer, Marten H. van Kerkwijk, Matthew Brett, Allan Haldane, Jaime Fernández del Río, Mark Wiebe, Pearu Peterson, Pierre Gérard-Marchant, Kevin Sheppard, Tyler Reddy, Warren Weckesser, Hameer Abbasi, Christoph Gohlke, and Travis E. Oliphant. Array programming with NumPy. Nature, 585(7825):357–362, September 2020. doi:10.1038/s41586-020-2649-2. URL https://doi.org/10.1038/s41586-020-2649-2.

[76] Richard J Gowers, Max Linke, Jonathan Barnoud, Tyler JE Reddy, Manuel N Melo, Sean L Seyler, Jan Domanski, David L Dotson, Sébastien Buchoux, Ian M Kenney, et al. MDAnalysis: a Python package for the rapid analysis of molecular dynamics simulations. In Proceedings of the 15th python in science conference, volume 98, page 105. SciPy Austin, TX, 2016.

[77] Naveen Michaud-Agrawal, Elizabeth J Denning, Thomas B Woolf, and Oliver Beckstein. MDAnalysis: a toolkit for the analysis of molecular dynamics simulations. Journal of computational chemistry, 32(10):2319–2327, 2011.

[78] J. D. Hunter. Matplotlib: A 2D graphics environment. Computing in Science & Engineering, 9(3):90–95, 2007. doi:10.1109/MCSE.2007.55.

[79] A.C.W. Pike, Y.Y. Dong, A. Tessitore, S. Goubin, C. Strain-Damerell, S. Mukhopadhyay, K. Kupinska, D. Wang, R. Chalk, G. Berridge, M. Grieben, L. Shrestha, J.H. Ang, A. Mackenzie, A. Quigley, S.R. Bushell, C.A. Shintre, B. Faust, A. Chu, L. Dong, F. von Delft, C.H. Arrowsmith, A.M. Edwards, C. Bountra, N.A. Burgess-Brown, and E.P. Carpenter. Crystal structure of human two pore domain potassium ion channel trek1 (k2p2.1), August 2014. URL 10.2210/pdb4twk/pdb.

[80] Wojciech Kopec, Brad S. Rothberg, and Bert L. de Groot. Molecular mechanism of a potassium channel gating through activation gate-selectivity filter coupling. Nature Communications, 10(1), November 2019. ISSN 2041-1723. doi:10.1038/s41467-019-13227-w. URL http://dx.doi.org/10.1038/s41467-019-13227-w.

[81] Pauli Virtanen, Ralf Gommers, Travis E. Oliphant, Matt Haberland, Tyler Reddy, David Cournapeau, Evgeni Burovski, Pearu Peterson, Warren Weckesser, Jonathan Bright, Stéfan J. van der Walt, Matthew Brett, Joshua Wilson, K. Jarrod Millman, Nikolay Mayorov, Andrew R. J. Nelson, Eric Jones, Robert Kern, Eric Larson,J J Carey, İlhan Polat, Yu Feng, Eric W. Moore, Jake VanderPlas, Denis Laxalde, Josef Perktold, Robert Cimrman, Ian Henriksen, E. A. Quintero, Charles R. Harris, Anne M. Archibald, Antônio H. Ribeiro, Fabian Pedregosa, Paul van Mulbregt, and SciPy 1.0 Contributors. SciPy 1.0: Fundamental Algorithms for Scientific Computing in Python. Nature Methods, 17:261–272, 2020. doi:10.1038/s41592-019-0686-2.

[82] Cédric Bouysset and Sébastien Fiorucci. Prolif: a library to encode molecular interactions as fingerprints. Journal of Cheminformatics, 13(1), September 2021. ISSN 1758-2946. doi:10.1186/s13321-021-00548-6. URL http://dx.doi.org/10.1186/s13321-021-00548-6.

[83] F. Pedregosa, G. Varoquaux, A. Gramfort, V. Michel, B. Thirion, O. Grisel, M. Blondel, P. Prettenhofer, R. Weiss, V. Dubourg, J. Vanderplas, A. Passos, D. Cournapeau, M. Brucher, M. Perrot, and E. Duchesnay. Scikit-learn: Machine learning in Python. Journal of Machine Learning Research, 12:2825–2830, 2011.

[84] Schrödinger, LLC. The PyMOL molecular graphics system, version 2.5.4. PyMOL, November 2015.

[85] William Humphrey, Andrew Dalke, and Klaus Schulten. VMD: visual molecular dynamics. Journal of molecular graphics, 14(1):33–38, 1996.

[86] Wojciech Kopec, Andrew S. Thomson, Bert L. de Groot, and Brad S. Rothberg. Interactions between selectivity filter and pore helix control filter gating in the mthk channel. Journal of General Physiology, 155(8), June 2023. ISSN 1540-7748. doi:10.1085/jgp.202213166. URL http://dx.doi.org/10.1085/jgp.202213166.

[87] Simone Furini and Carmen Domene. Critical assessment of common force fields for molecular dynamics simulations of potassium channels. Journal of Chemical Theory and Computation, 16(11):7148–7159, October 2020. ISSN 1549-9626. doi:10.1021/acs.jctc.0c00331. URL http://dx.doi.org/10.1021/acs.jctc.0c00331.

[88] Victoria Oakes, Simone Furini, David Pryde, and Carmen Domene. Exploring the dynamics of the twik-1 channel. Biophysical Journal, 111(4):775–784, August 2016. ISSN 0006-3495. doi:10.1016/j.bpj.2016.07.009. URL http://dx.doi.org/10.1016/j.bpj.2016.07.009.

[89] Carus H.Y. Lau, Emelie Flood, Mark J. Hunter, Billy J Williams-Noonan, Karen M. Corbett, Chai-Ann Ng, James C. Bouwer, Alastair G. Stewart, Eduardo Perozo, Toby W. Allen, and Jamie I. Vandenberg. Potassium dependent structural changes in the selectivity filter of herg potassium channels. bioRxiv, 2023. doi:10.1101/2023.12.14.571769. URL https://www.biorxiv.org/content/early/2023/12/15/2023.12.14.571769.

[90] E Nicholas Petersen, Mahmud Arif Pavel, Samuel S Hansen, Manasa Gudheti, Hao Wang, Zixuan Yuan, Keith R Murphy, William Ja, Heather A Ferris, Erik Jorgensen, et al. Mechanical activation of twik-related potassium channel by nanoscopic movement and rapid second messenger signaling. Elife, 12, December 2024. doi:10.7554/elife.89465.2. URL http://dx.doi.org/10.7554/eLife.89465.2.

[91] Sviatoslav N Bagriantsev, Kimberly A Clark, and Daniel L Minor. Metabolic and thermal stimuli control K(2P)2.1 (TREK-1) through modular sensory and gating domains. The EMBO Journal, 31(15):3297–3308, June 2012. ISSN 0261-4189. doi:10.1038/emboj.2012.171. URL http://dx.doi.org/10.1038/emboj.2012.171.

[92] Karin E.J. Rödström, Alexander Cloake, Janina Sörmann, Agnese Baronina, Kathryn H.M. Smith, Ashley C.W. Pike, Jackie Ang, Peter Proks, Marcus Schewe, Ingelise Holland-Kaye, Simon R. Bushell, Jenna Elliott, Els Pardon, Thomas Baukrowitz, Raymond J. Owens, Simon Newstead, Jan Steyaert, Elisabeth P. Carpenter, and Stephen J. Tucker. Extracellular modulation of trek-2 activity with nanobodies provides insight into the mechanisms of k2p channel regulation. bioRxiv, 2023. doi:10.1101/2023.10.19.562917. URL https://www.biorxiv.org/content/early/2023/12/12/2023.10.19.562917.

[93] Yunguang Qiu, Lu Huang, Jie Fu, Chenxia Han, Jing Fang, Ping Liao, Zhuo Chen, Yiqing Mo, Peihua Sun, Daqing Liao, Linghui Yang, Jing Wang, Qiansen Zhang, Jin Liu, Feng Liu, Tingting Liu, Wei Huang, Huaiyu Yang, and Ruotian Jiang. TREK Channel Family Activator with a Well-Defined Structure–Activation Relationship for Pain and Neurogenic Inflammation. Journal of Medicinal Chemistry, 63(7):3665–3677, March 2020. ISSN 1520-4804. doi:10.1021/acs.jmedchem.9b02163. URL http://dx.doi.org/10.1021/acs.jmedchem.9b02163.

